# *RPS19* and *RPL5*, the most commonly mutated genes in Diamond Blackfan anemia, impact DNA double-strand break repair

**DOI:** 10.1101/2024.10.10.617668

**Authors:** Nicholas F. DeCleene, Elif Asik, Anthony Sanchez, Christopher L. Williams, Elena B. Kabotyanski, Na Zhao, Nimrat Chatterjee, Kyle M. Miller, Yu-Hsiu Wang, Alison A. Bertuch

## Abstract

Diamond Blackfan anemia (DBA) is caused by germline heterozygous loss-of-function pathogenic variants (PVs) in ribosomal protein (RP) genes, most commonly *RPS19* and *RPL5*. In addition to red cell aplasia, individuals with DBA are at increased risk of various cancers. Importantly, the mechanism(s) underlying cancer predisposition are poorly understood. We found that DBA patient-derived lymphoblastoid cells had persistent γ-H2AX foci following ionizing radiation (IR) treatment, suggesting DNA double-strand break (DSB) repair defects. RPS19- and RPL5-knocked down (KD) CD34+ cells had delayed repair of IR-induced DSBs, further implicating these RPs in DSB repair. Assessing the impact of RPS19- and RPL5-KD on specific DSB repair pathways, we found RPS19-KD decreased the efficiency of pathways requiring extensive end-resection, whereas RPL5-KD increased end-joining pathways. Additionally, RAD51 was reduced in RPS19- and RPL5-KD and RPS19- and RPL5-mutated DBA cells, whereas RPS19-deficient cells also had a reduction in PARP1 and BRCA2 proteins. RPS19-KD cells had an increase in nuclear RPA2 and a decrease in nuclear RAD51 foci post-IR, reflective of alterations in early, critical steps of homologous recombination. Notably, RPS19 and RPL5 interacted with poly(ADP)-ribose chains noncovalently, were recruited to DSBs in a poly(ADP)-ribose polymerase activity-dependent manner, and interacted with Ku70 and histone H2A. RPL5’s recruitment, but not RPS19’s, also required p53, suggesting that RPS19 and RPL5 directly participate in DSB repair via different pathways. We propose that defective DSB repair arising from haploinsufficiency of these RPs may underline the cancer predisposition in DBA.

**KEY POINTS:**

- RPS19- and RPL5-deficiency alters DNA double-strand break repair, which may underlie cancer predisposition in Diamond Blackfan anemia.
- RPS19 and RPL5 bind PAR chains and are rapidly recruited to sites of DNA double-strand breaks, suggesting active roles in repair.

## INTRODUCTION

Diamond Blackfan anemia (DBA) is a rare inherited bone marrow failure syndrome most often caused by germline heterozygous loss-of-function mutations in one of 24 ribosomal protein (RP) genes.^1,2^ Studies have pointed to different mechanisms underlying anemia in DBA.^3^ For example, reduced numbers of ribosomes alter the translation efficiency (TE) of the GATA1 transcript, decreasing protein expression of this master transcriptional regulator of erythropoiesis.^4^ In addition, dysregulated ribosome biogenesis results in the binding of the RPL5/RPL11/5S ribosomal RNA complex, called the 5S ribonuclear protein (RNP), or other free RPs to mouse double minute 2 (MDM2), competitively inhibiting its ubiquitination of p53, leading to p53 stabilization and cell cycle arrest of erythroid progenitors.^5–7^

DBA is also a cancer predisposition syndrome, presenting an increased risk of myelodysplastic syndrome, acute myeloid leukemia, and a range of solid tumors.^8,9^ In particular, there is a 40-fold increased risk of colorectal cancer, which is also early onset, and osteosarcoma compared to the general population. Importantly, the mechanism(s) underlying cancer predisposition in DBA, while of fundamental biological and clinical interest, remain poorly understood.^10^

In addition to serving as ribosome components, several RPs have extraribosomal functions, including in DNA repair. For example, RPS3 contributes endonuclease activity to base and nucleotide excision repair of ultraviolet-induced DNA damage and functions as a negative regulator of the nonhomologous end-joining (NHEJ) pathway of DNA double-strand break (DSB) repair.^11–14^ RPL6 is recruited to DSBs, where it modulates the DNA damage response (DDR) through interactions with histone H2A and negatively regulates NHEJ and homologous recombination (HR) pathways.^15^ To date, none of the DBA-associated RPs has been reported to be involved in DNA repair.

There are four DSB repair pathways: NHEJ, alternative end-joining (alt-EJ), single-strand annealing (SSA), and HR. The first three are mutagenic, whereas HR is a high-fidelity process that utilizes a homologous template for repair.^16^ In this study, we uncover differential influences of the two most frequently mutated RPs in DBA, RPS19 and RPL5, on DSB repair and present evidence that supports extraribosomal roles of these RPs in the repair of DSBs. Our data suggest that haploinsufficiency of RPS19 or RPL5 shifts DSB repair away from high-fidelity and towards error-prone repair, which may contribute to cancer predisposition in DBA.

## METHODS

### Immunofluorescence (IF) assays

IF assays were performed using standard methods (see supplemental methods).

### Neutral comet assays

Neutral comet assays were conducted according to the Bio-Techne protocol (4250-050-5) using CD34+ cells following transfection of siControl (Scr), RPS19, and RPL5 siRNAs.

### HR, total-EJ, SSA, and alt-EJ reporter assays

DSB repair pathway assays were performed using U2OS cells with integrated previously described reporters.^17–19^ Repair efficiency was calculated by the percent of GFP to mCherry (transfection control) positive cells.

### Polysome profiling

Polysome profiling experiments were conducted as described previously^20^ and are detailed further in the supplemental methods.

### Laser microirradiation

Laser microirradiation experiments were conducted as described previously^21^ and are detailed further in the supplemental methods.

### FokI co-localization assays

FokI assays were conducted using reporter cell lines as described previously.^22^ Site-specific DSBs were induced by adding 1 µM Shield-1 (Takara, 632189) and 1 µM 4-hydroxytamoxifen (Sigma, H7904-5MG) for 5 hours before fixation. Cells were fixed as described previously.^23^ Detailed information regarding imaging conditions can be found in supplemental methods.

### Full Methodology

Detailed methods are provided in Supplemental Information.

### Data Sharing Policy

For original data, please contact.

## RESULTS

### Lymphoblastoid cells derived from individuals with DBA and RPS19- and RPL5-depleted CD34+ cells exhibit impaired DSB repair

To investigate if RPS19 and RPL5, the two most commonly mutated genes implicated in DBA, contribute to DSB repair, we tested Epstein Barr virus (EBV)-transformed lymphoblastoid cell lines (LCLs) derived from individuals with germline *RPS19* and *RPL5* PVs (supplemental Table 1) for increased DNA damage following IR. Compared to pooled healthy control LCLs, each DBA LCL had more DNA damage as assayed by the DNA damage marker γ-H2AX before and 24 hours following treatment (Figure 1A–B). To determine if this phenotype, which was suggestive of a defect in DSB repair, was secondary to EBV transformation, we examined the resolution kinetics of DSBs induced by IR in RPS19- and RPL5-siRNA knocked down (KD) bone marrow CD34+ cells by neutral comet assays.^24^ We found both RPS19- and RPL5-KD CD34+ cells had delayed resolution of IR-induced DSBs compared to control cells, with persistently increased tail moments (i.e., DSBs) 24 hours post-IR (Figure 1C–H; supplemental Figure 1A–B). Similar results were observed using multiple siRNAs targeting RPS19 and RPL5 (supplemental Table 2), arguing against off-target effects (supplemental Figure 1C). These data demonstrate that RPS19 and RPL5 influence the repair of DSBs.

**Figure 1.**
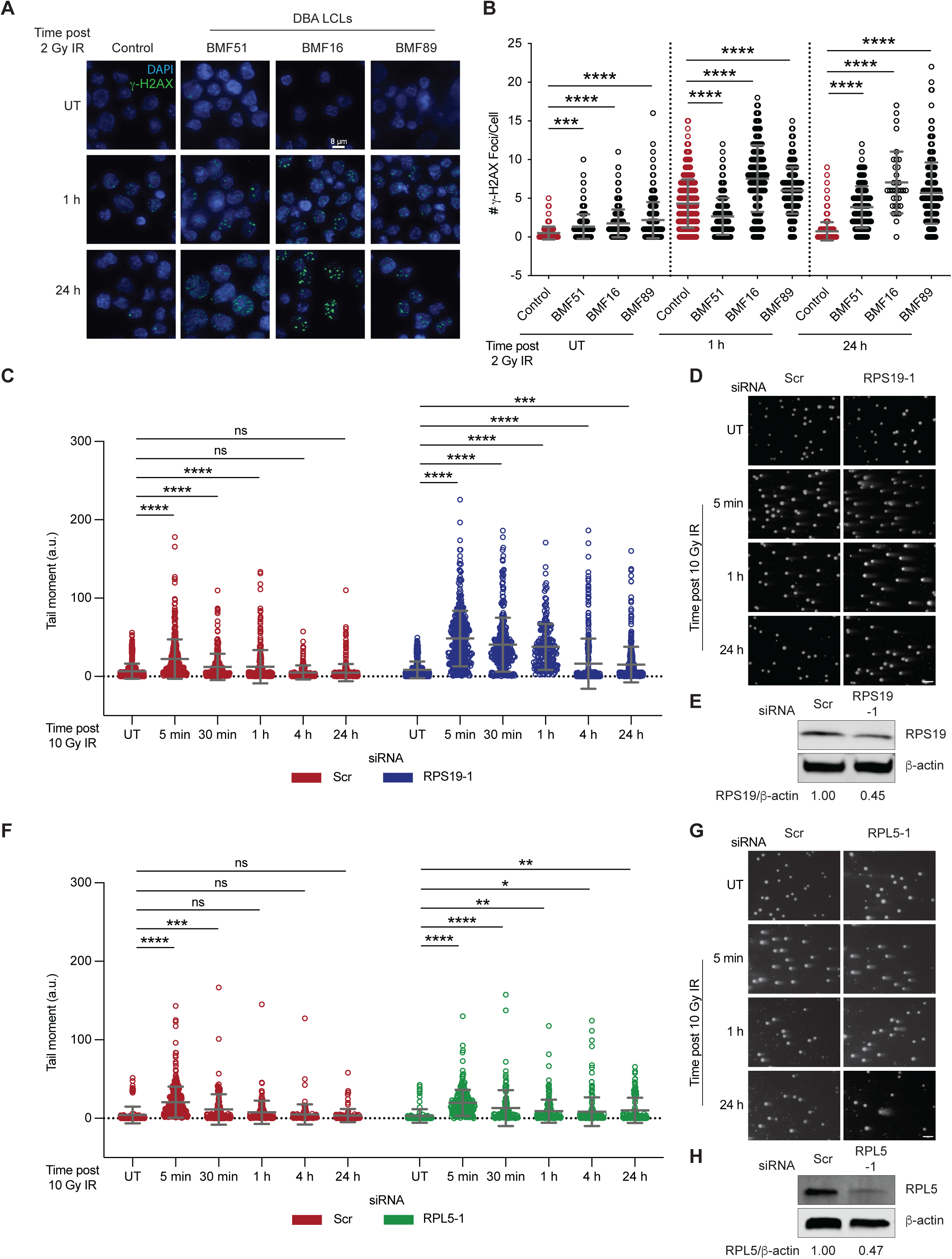
DBA patient-derived LCLs with mutations in *RPS19* or *RPL5* have delayed resolution of γ-H2AX foci and RPS19- and RPL5-knocked down (KD) CD34+ cells have impaired DSB repair. (A) Representative images of cells analyzed for γ-H2AX foci from DBA or a control LCL that were untreated (UT) or treated with 2 Gy IR and harvested at 1 and 24 hr. DAPI was used for nuclear counterstaining. Bar, 8 µm. (B) Quantification of γ-H2AX foci in LCLs treated as in A using high throughput microscopy and single-cell image analysis. A minimum of 34 cells were analyzed per condition. (C) Neutral comet assays of siRNA KD RPS19-1 or scrambled (Scr) CD34+ cells that were UT or treated with 10 Gy IR were analyzed at the indicated time points post IR. (D) Representative images of cells analyzed in C. Bar, 50 µM. (E) Western blot of RPS19-1 KD CD34+ cells assessing the protein levels of RPS19. β-actin is used as a loading control. (F) Neutral comet assays of siRNA KD RPL5-1 or Scr CD34+ cells UT or treated with 10 Gy IR and analyzed at the indicated time points post IR. (G) Representative images of cells analyzed in F. Bar, 50 µM. (H) Western blot of RPL5-1 KD CD34+ cells assessing the protein levels of RPL5. β-actin as a loading control. A minimum of 115 cells were analyzed per condition. Data represent mean ± SD compared using 1-way ANOVA with Dunnett’s multiple comparisons test. Ns not significant; \**P* < 0.05; \*\**P* < 0.01; \*\*\**P* < 0.001; \*\*\*\**P* < 0.0001.

### Decreased HR repair in RPS19-KD cells and increased total-EJ repair in RPL5-KD cells

Given the delay in DSB repair, we wondered if specific DSB repair pathways were impacted by RPS19 or RPL5 deficiency. We utilized well-established I-SceI/GFP-based DSB repair reporter systems to assess the impact of RPS19- and RPL5-KD on the efficiency of total end-joining (total-EJ), comprising error-free or mutagenic NHEJ and alt-EJ, and HR in U2OS cells.^16^ For each protein, we tested the two unique siRNAs (supplemental Table 2) to validate effects (Figure 2A), and a mCherry-expression plasmid as a transfection control. RPS19-KD resulted in a 50% reduction in HR but no change in total-EJ (Figure 2B–C). In addition to observing similar effects with the two distinct siRNAs, we found that ectopic expression of siRNA-resistant and flag-tagged RPS19 in RPS19-deficient cells restored the HR efficiency, confirming an on-target effect (Figure 2D–E). We also tested RPS19-KD HCT116 cells and found similar HR and total-EJ results (supplemental Figure 2A–B). Together, these results indicate that efficient DSB repair via HR is sensitive to reduced RPS19.

**Figure 2.**
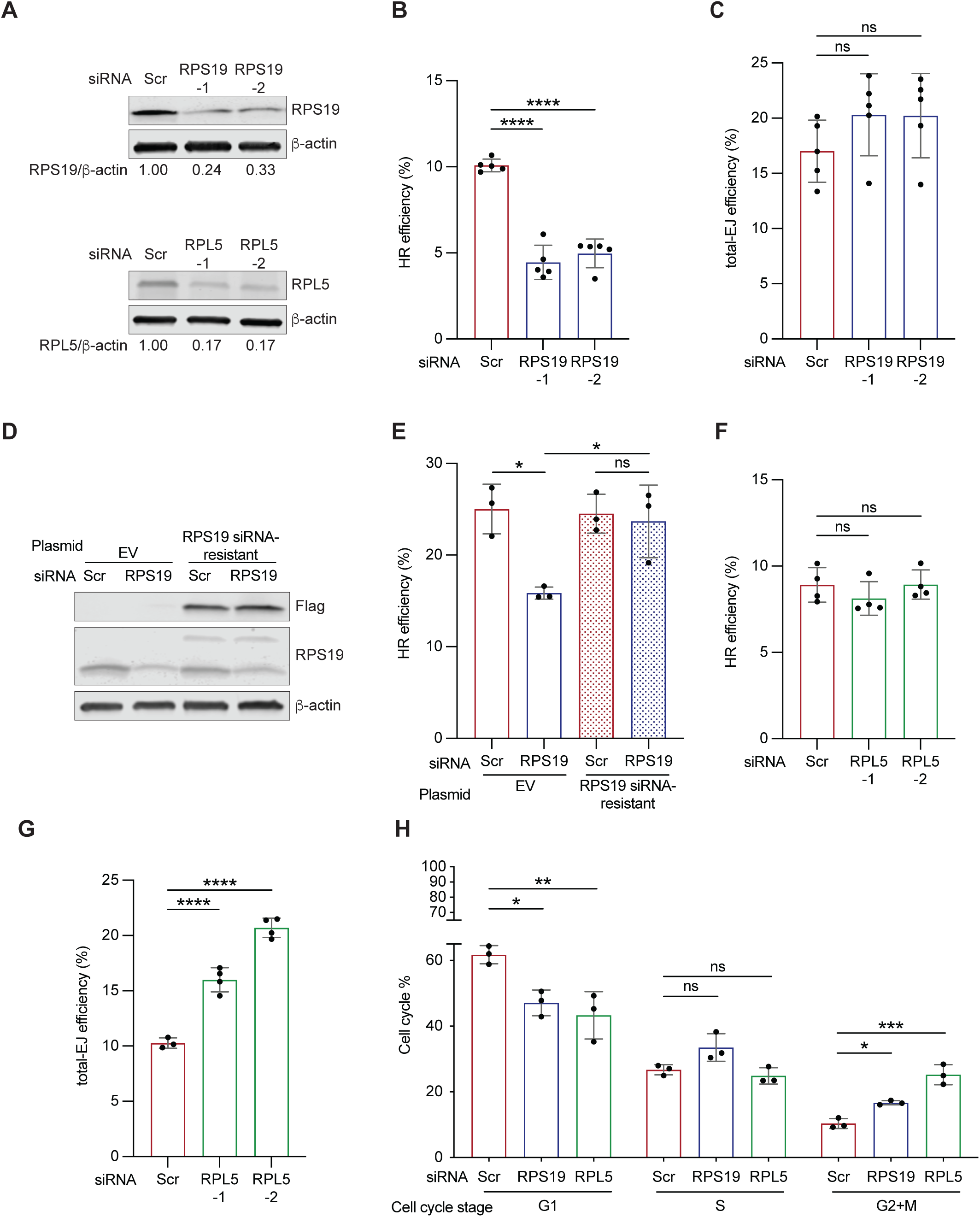
RPS19-KD U2OS cells have decreased HR repair, whereas RPL5-KD U2OS cells have increased total-EJ repair. (A) Western blots of RPS19 and RPL5 KD U2OS cells using two different siRNAs for each (RPS19-1, RPS19-2, RPL5-1, and RPL5-2) that target different regions of each transcript. Probed for RPS19 or RPL5. β-actin is used as a loading control. (B) U2OS cells bearing an integrated DR-GFP HR reporter^17^ transfected with Scr or one of two RPS19 siRNAs. After 48 hours, cells were co-transfected with an I-SceI-expressing plasmid to induce a DSB in the DR-GFP reporter and a mCherry-expressing plasmid as a transfection control. Forty-eight hours later, cells were analyzed by flow cytometry. The repair efficiency was calculated by the proportion of GFP+ to mCherry+ cells. (C) The same experiment as in B except using U2OS cells with an integrated EJ5-GFP total-EJ reporter.^18^ (D) Western blot of U2OS cells with RPS19-1 KD and overexpression of an RPS19 siRNA-resistant plasmid with a 3x-FLAG tag. Probed for RPS19 and FLAG. β-actin used as a loading control. (E) The same experiment as in B, except for an RPS19 siRNA-resistant or an empty vector (EV) plasmid was transfected along with RPS19-1 or Scr siRNA. (F) The same experiment as in B except for using two different RPL5 siRNAs. (G) The same experiment as in C except for using two different RPL5 siRNAs. (H) U2OS cells transfected with Scr, RPS19-1, or RPL5-1 siRNAs for 72 hours, followed by flow cytometry using 7-AAD and BrdU. Data represent mean ± SD compared using 1-way ANOVA with Dunnett’s multiple comparisons test of a minimum of three biological replicates. ns not significant; \**P* < 0.05; \*\**P* < 0.01; \*\*\**P* < 0.001; \*\*\*\**P* < 0.0001.

Surprisingly, when we assessed the impact of RPL5-KD on HR and total-EJ, we found different results: HR efficiency was unaffected, whereas the efficiency of total-EJ was increased (Figure 2F–G). The opposing effects extended to the two less frequently utilized DSB repair pathways, SSA and alt-EJ. Using SSA and alt-EJ I-SceI/GFP-based reporter cell lines,^18,19^ we found that RPS19-KD U2OS cells had decreased SSA and unchanged alt-EJ, whereas RPL5-deficient cells had unchanged SSA and increased alt-EJ (supplemental Figure 2C–F). Since total-EJ assesses both alt-EJ and NHEJ repair events, we compared the ratio of alt-EJ to total-EJ repair events as described previously.^25^ We found that RPL5-1 and RPL5-2 KD cells had a relative foldchange of 1.1 and 1.3, respectively. These data suggest that increased alt-EJ drives the increase in total-EJ. These data indicate that RPS19 promotes DSB repair pathways that utilize extensive end resection, whereas RPL5 suppresses end-joining pathways.

### Changes in cell cycle profiles do not account for altered DSB repair pathway efficiencies in RPS19- and RPL5-KD cells

The utilization of HR and NHEJ varies across the cell cycle: HR occurs mainly during late S and G2 when a sister chromatid is available to serve as a template for repair, whereas end-joining events occur throughout the cell cycle, primarily in G1.^26^ To assess if the alterations in DSB repair upon RPS19- or RPL5-KD could be due to changes in cell cycle progression, we performed cell cycle analysis using BrdU labeling and flow cytometry. RPS19- and RPL5-KD cells had a decrease in the proportion of cells in G1, no change in the proportion in S phase, and an increase in the proportion in G2/M, which suggests unrepaired damage (Figure 2H). Given that RPS19-KD cells had decreased HR, but the proportion of cells was unchanged or increased in the S and G2/M stages of the cell cycle, respectively, a reduction in HR could not be attributed to fewer cells in the S and G2 stages. Furthermore, RPL5-KD cells had increased total-EJ while the proportion of cells in G1 was decreased; thus, the increase in total-EJ is unlikely to be accounted for by an increase in cells in G1. RPS19- and RPL5-KD cells demonstrated decreased proliferation rates (supplemental Figure 2G), which could be related to the altered cell cycle distributions. However, the similarity suggests that the decrease does not underlie the differential impacts on DSB repair.

### RPS19-KD cells have decreased RAD51, BRCA2, and PARP1 protein levels

Next, we investigated if the changes in the efficiencies of specific DSB repair pathways in RPS19- and RPL5-KD cells could be due to changes in the level of one or more pathway-specific proteins. We analyzed the levels of over 20 targets with representative key repair factors from 5 different repair pathways. Whereas KD of RPS19 or RPL5 did not cause a global reduction of protein abundance (Figure 3A–B; supplemental Figure 3 and 4), they led to a decrease in RAD51 levels. This decrease was more pronounced in RPS19-KD cells (with a mean of 20-40% RAD51 remaining), which exhibited decreased HR efficiency (Figure 2B), compared to RPL5-KD cells (mean of 50-60% RAD51 remaining), where HR was preserved (Figure 2G). Thus, the influence of RPS19 on HR efficiency could be due, at least in part, to RAD51 levels falling below the threshold necessary to maintain normal efficiency. A previous report noted that RAD51 is a common off-target in siRNA studies.^27^ Corroborating that RPS19 or RPL5 deficiency leads to a reduction in RAD51, we found the level of RAD51 was reduced in LCLs derived from individuals with mutated *RPS19*- or *RPL5* DBA compared to the level in LCLs derived from healthy donors (Figure 3C). This rules out off-target siRNA effects and supports that RPS19 and RPL5 impact RAD51 levels.

**Figure 3.**
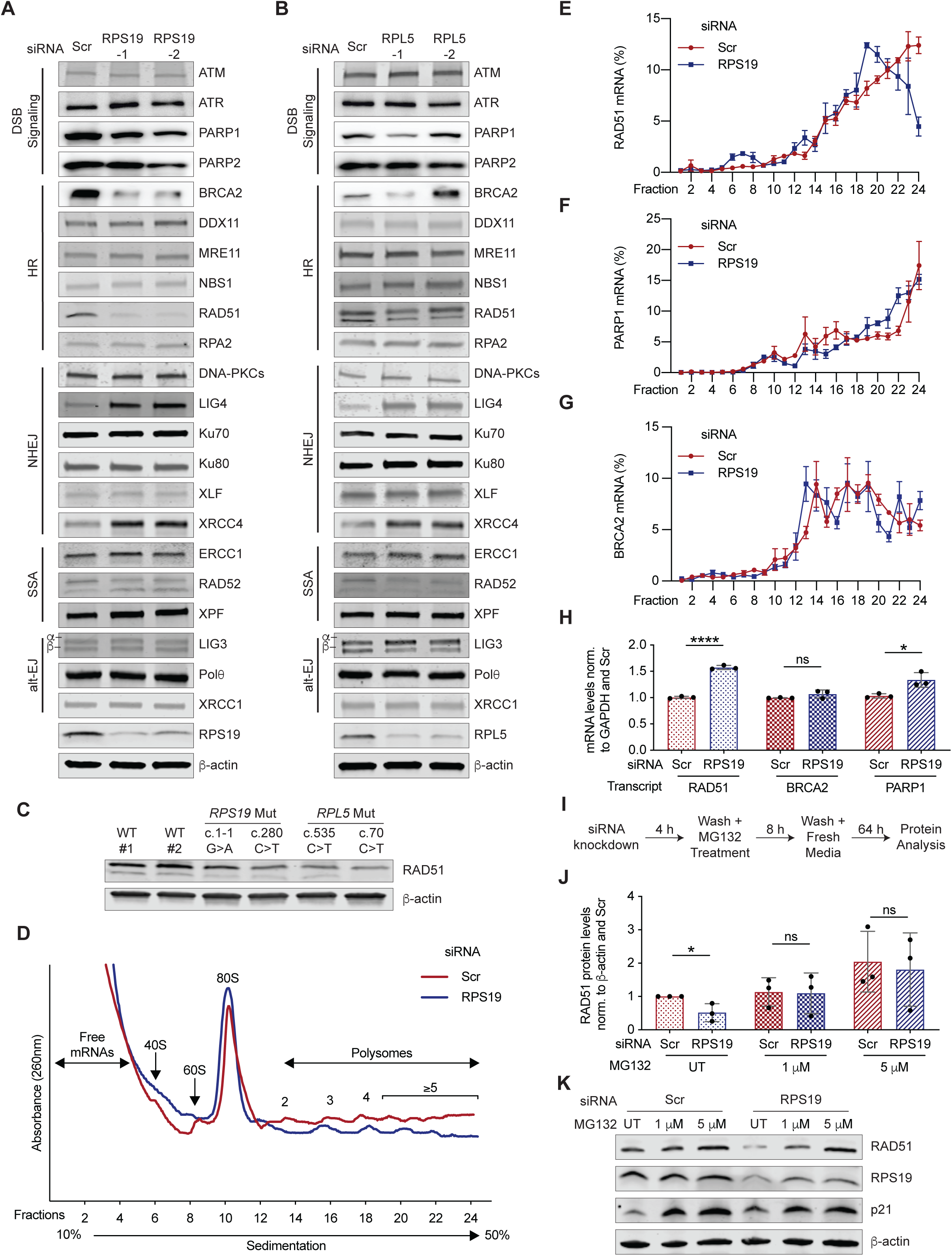
RPS19-KD U2OS cells have decreased RAD51, BRCA2, and PARP1 protein levels and RAD51 mRNA has reduced association with polysomes in RPS19-KD U2OS cells. (A) Western blots of RPS19-KD U2OS cells assessing the protein levels of DSB repair factors. β- actin is used as a loading control. (B) Western blots of RPL5-KD U2OS cells assessing the protein levels of DSB repair factors. β-actin is used as a loading control. RAD51 has been shown to have one or two bands depending on the gel percentage used. In both instances, siRNA knockdown has validated the RAD51 band. (C) Western blot assessing RAD51 levels in DBA LCLs derived from healthy control individuals and individuals with pathogenic RPS19 or RPL5 variants. β-actin is used as a loading control. (D) Representative polysome profiles of Scr and RPS19-1 KD U2OS cells. (E) Distribution of RAD51 mRNAs across the different fractions in Scr and RPS19-1 KD U2OS cells. (F) Same as E except assessing PARP1 mRNAs. (G) Same as E except assessing BRCA2 mRNAs. (H) Quantification of RAD51, BRCA2, and PARP1 mRNA levels in RPS19-1 KD U2OS cells normalized to GAPDH and Scr. (I) Schema of proteasomal inhibition experiment. (J) Quantification of RAD51 protein levels in RPS19-1 KD U2OS cells normalized to β-actin and Scr. p21 is used as positive control for proteasome inhibition. (K) Western blot of lysates prepared from RPS19-1 KD cells treated with the proteasomal inhibitor MG132. β-actin is used as a loading control. Data in E–G represent the mean ± SEM of three biological replicates. Data in H and J represent mean ± SD compared using unpaired t-tests of three biological replicates. ns not significant; \**P* < 0.05; \*\*\*\**P* < 0.000.

We also found reduced levels of BRCA2, which is a key HR factor, and PARP1, which negatively regulates end resection, in RPS19-KD cells. Lastly, the levels of ERCC1, RAD52, and XPF, which are required for SSA, were not appreciably reduced in RPS19-KD cells, yet SSA was reduced (Figure 3A; supplemental Figure 2C and 3E), suggesting pathways other than reduced protein levels may influence DSB repair in RPS19-KD cells.

In their previous studies of RPS19- and RPL5-depleted primary hematopoietic cells undergoing erythroid differentiation, Khajuria et al.^4^ identified a selective reduction in the TE. However, neither RAD51, BRCA2, nor PARP1 were identified among the transcripts sensitive to ribosome reduction. To test whether translation was affected, we performed polysome profiling and targeted mRNA analysis of these transcripts in RPS19-KD and Scr control cells (Figure 3D-G). We found a small peak of RAD51 transcript preceding the 80S ribosomal peak fraction in the RPS19-KD samples, which was absent in the Scr samples. The remaining RAD51 mRNA was distributed toward the lighter polysome fractions in the RPS19-KD samples, peaking around 4-mer polysomes in contrast to Scr samples, which peaked at >5-mer polysomes (Figure 3E). This shift in RAD51’s association with lighter polysomes was further illustrated by the greater number of combined heaviest to lightest fractions to reach 50% of the total RAD51 mRNA (supplemental Figure 5A). In contrast, we found an increase in the association of PARP1 mRNA with heavier polysomes and no change in BRCA2 mRNA in the RPS19-KD cells compared to the Scr cells (Figure 3F–G; supplemental Figure 5B–C). Separately, we analyzed the bulk mRNA levels and found a modest increase in RAD51, which could offset the decrease in TE, no change in BRCA2, and a smaller increase in PARP1 mRNA in RPS19-KD cells (Figure 3H). Lastly, we examined the contribution of the proteasome, focusing on RAD51. We found that treating RPS19-KD cells with the proteasome inhibitor MG132 restored the RAD51 protein level to the level of comparably treated Scr control cells (Figure 3I-K). Thus, proteasomal degradation influences the level of RAD51 in RPS19-KD cells, which could contribute to the HR defect.

### RPS19- and RPL5-KD cells have increased levels of the NHEJ factors LIG4 and XRCC4

In addition, NHEJ factors LIG4 and XRCC4 were markedly increased in RPS19- and RPL5-deficient cells (Figure 3A–B; supplemental Figure 3D and 4D). LIG4 mRNA was similarly increased in these cells, whereas XRCC4 mRNA was increased only in the RPS19-KD cells (supplemental Figure 5E–F). LIG4 and XRCC4 operate in a complex that catalyzes the ligation of DSB ends in NHEJ.^28^ Although the ratio of alt-EJ to total-EJ repair events in RPL5-depleted cells suggested alt-EJ rather than NHEJ was increased, it was important to determine whether the increased levels of LIG4 and XRCC4 contributed to the increased in total-EJ in RPL5-KD cells or the decreased HR repair in RPS19-KD cells. Previous studies showed that a reduction in XRCC4 accompanies LIG4-KD.^29^ Therefore, we used LIG4 siRNA to restore both proteins to relatively normal levels in RPS19- or RPL5-KD total-EJ and HR reporter cells and tested the repair efficiencies (supplemental Figure 6A–C). Despite reducing LIG4 and XRCC4 levels to around that of the Scr siRNA control cells, the total-EJ efficiency of RPL5 KD cells remained increased following LIG4 siRNA treatment. LIG4-KD did not impact total-EJ efficiency in the Scr control cells, suggesting that LIG4 was not reduced enough to impact NHEJ levels.^30^ Still, HR efficiency increased from 14% to 22%, consistent with previous findings.^29^ LIG4 depletion had a minor impact on HR in RPS19-KD cells, whereas it substantially increased it in RPL5-KD cells, suggesting other factors restrain HR when RPS19 is deficient. We also over-expressed LIG4 and XRCC4 in the HR and total-EJ reporter cells and found that they did not impact either pathway (supplemental Figure 6D–F). These data suggest that RPS19 does not promote HR, and RPL5 does not restrain total-EJ by limiting LIG4/XRCC4 protein levels.

### RPS19-KD cells have increased nuclear RPA2 and a lack of accumulation of RAD51 foci following DNA damage

Since the two pathways that were reduced in RPS19-KD cells, HR and SSA (Figure 2B; supplemental Figure 2C), require DNA-end resection, we examined whether end resection was restrained in these cells. RPA rapidly associates with single-stranded DNA (ssDNA) created upon resection of the 5’ strands at DSB ends, and, as such, it can be utilized as a surrogate for end resection.^31^ We treated RPS19-KD cells with 10 Gy of IR and assessed nuclear RPA2, a component of the RPA heterotrimer. We found that, before (UT) and at 30 minutes and 1 hour post IR treatment, RPS19-KD cells had greater nuclear RPA2 compared to Scr siRNA-treated control cells (Figure 4A–B) despite relatively stable total RPA protein levels (Figure 4C). These data indicate that the reduction in HR and SSA efficiency in RPS19-KD cells was not due to an inhibition of end resection. Notably, PARP1 was significantly reduced by RPS19-KD (Figure 3A; supplemental Figure 3B). As PARP1 is a negative regulator of end resection,^32^ its reduction may have contributed to the increase in RPA2 nuclear accumulation in RPS19-KD cells.

**Figure 4.**
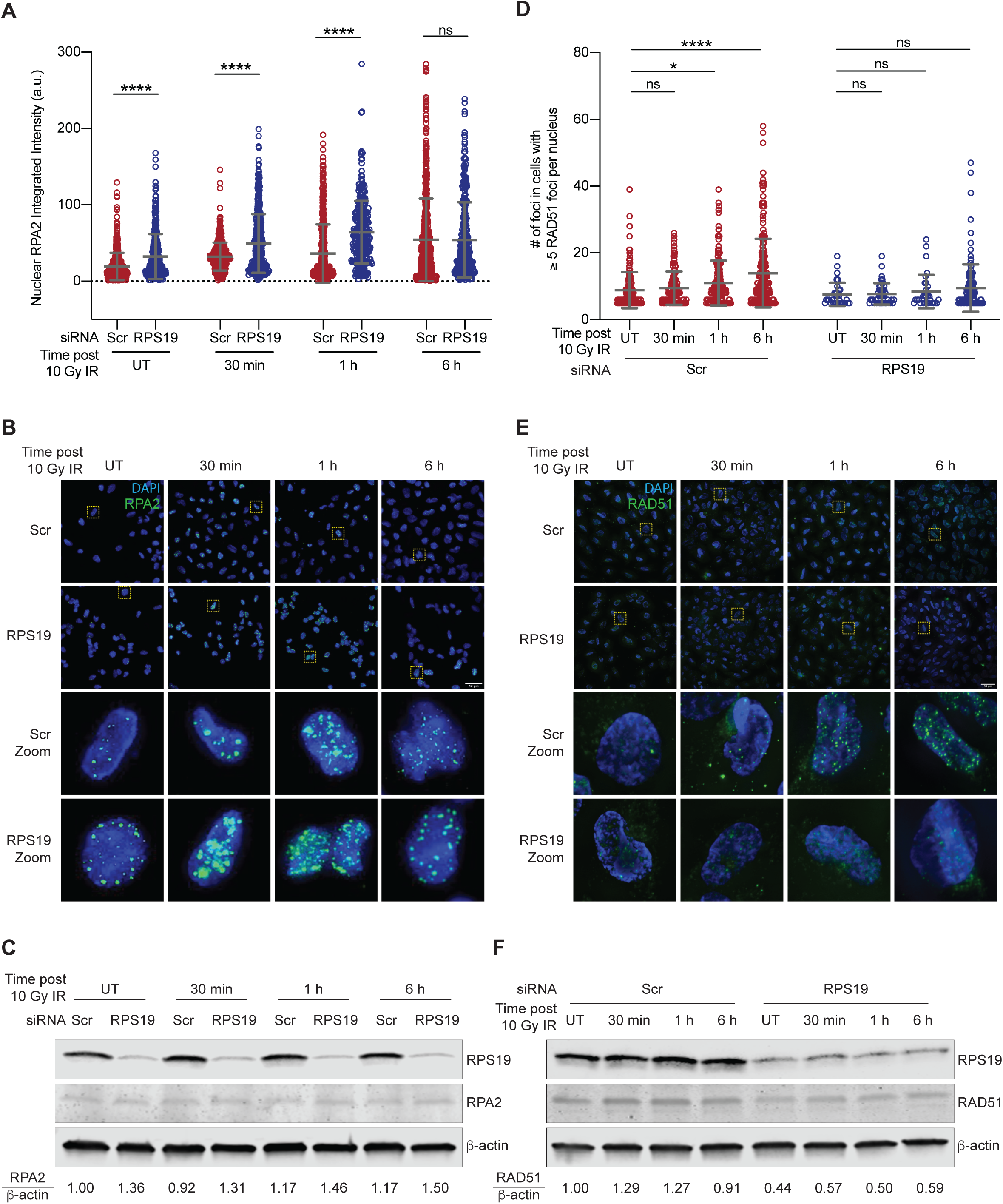
RPS19-KD cells have increased nuclear RPA2 integrated intensity and decreased nuclear RAD51 foci. (A) Quantification of nuclear RPA2 integrated intensity in RPS19-1 KD U2OS cells pre-treatment (UT) and following treatment with 10 Gy IR. Representative data of two biological replicates. (B) Representative images of nuclear RPA2 (green) analyzed in A. DAPI (blue) was used for nuclear counterstaining. Yellow box denotes zoomed area. (C) Western blot of total RPA2 protein levels in RPS19-1 KD U2OS cells represented in A. β-actin used as a loading control. (D) Quantification of cells with ≥5 nuclear RAD51 foci in RPS19-1 KD U2OS cells pre-treatment (UT) and following treatment with 10 Gy IR. Representative data of two biological replicates which were combined. (E) Representative images of nuclear RAD51 foci (green) analyzed in D. DAPI (blue) was used for nuclear counterstaining. Yellow box denotes zoomed area. (F) Western blot of total RAD51 protein levels in RPS19-1 KD U2OS cells represented in A. β-actin used as a loading control. Data in A represent mean ± SD compared using unpaired t-tests. Data in D represent mean ± SD compared using 1-way ANOVA with Dunnett’s multiple comparisons test. ns not significant; \**P* < 0.05; \*\*\*\**P* < 0.0001

Next, we investigated whether a step downstream of DNA end-resection might be compromised. Following RPA’s loading onto ssDNA, BRCA2 mediates the nucleation of RAD51 filaments,^33,34^ a critical step for subsequent homology search and strand invasion. Consistent with impairment of this exchange, while some RAD51 were detected, the number of RAD51 foci remained unchanged in RPS19-KD cells following treatment in contrast to the increase in RAD51 foci numbers observed in the Scr control cells at 1 hour and 6 hours post IR treatment (Figure 4D–F). Taken together, the increase in nuclear RPA2, the decrease in BRCA2 protein, and the lack of increase in RAD51 foci in RPS19-KD cells suggest that not only is RAD51 protein reduced, but the exchange of RPA2 with RAD51 is perturbed, and this could be contributing to the observed defects in HR.

### Fluorescently tagged RPS19 and RPL5 are rapidly recruited to sites of DSBs in a PARP-dependent and ATM-influenced manner, and their localization at DSBs requires noncovalent PAR binding

To examine if RPS19 or RPL5 has a more direct role in DSB repair, we examined their recruitment to sites of DSBs. First, we analyzed eGFP-RPS19 and mCherry-RPL5 recruitment to sites of laser microirradiation using live cell imaging. We found that both proteins were rapidly recruited to DNA lesions within 30 seconds of laser-induced damage (Figure 5A; supplemental Figure 7A). The recruitment of RPS19 was impaired by the PARP inhibitor veliparib and enhanced by the PARG inhibitor PDD 00017273 (Figure 5B). RPL5’s recruitment was also inhibited by veliparib, but in contrast to RPS19’s recruitment, it was delayed and blunted by PARG inhibition (Figure 5C). To corroborate these findings, we utilized a U2OS cell line expressing a mCherry-LacI-FokI nuclease and analyzed the co-localization of eGFP-RPS19 and eGFP-RPL5 with the nuclease-generated DSBs.^22^ Both eGFP-RPS19 and eGFP-RPL5 localized to FokI sites, and their recruitment diminished in the presence of the PARP inhibitor olaparib, consistent with the laser microirradiation studies (Figure 5D–E). These data demonstrate that both proteins accumulate within the vicinity of DSBs and support a model in which RPS19 and RPL5 play an active role in DSB repair.

**Figure 5.**
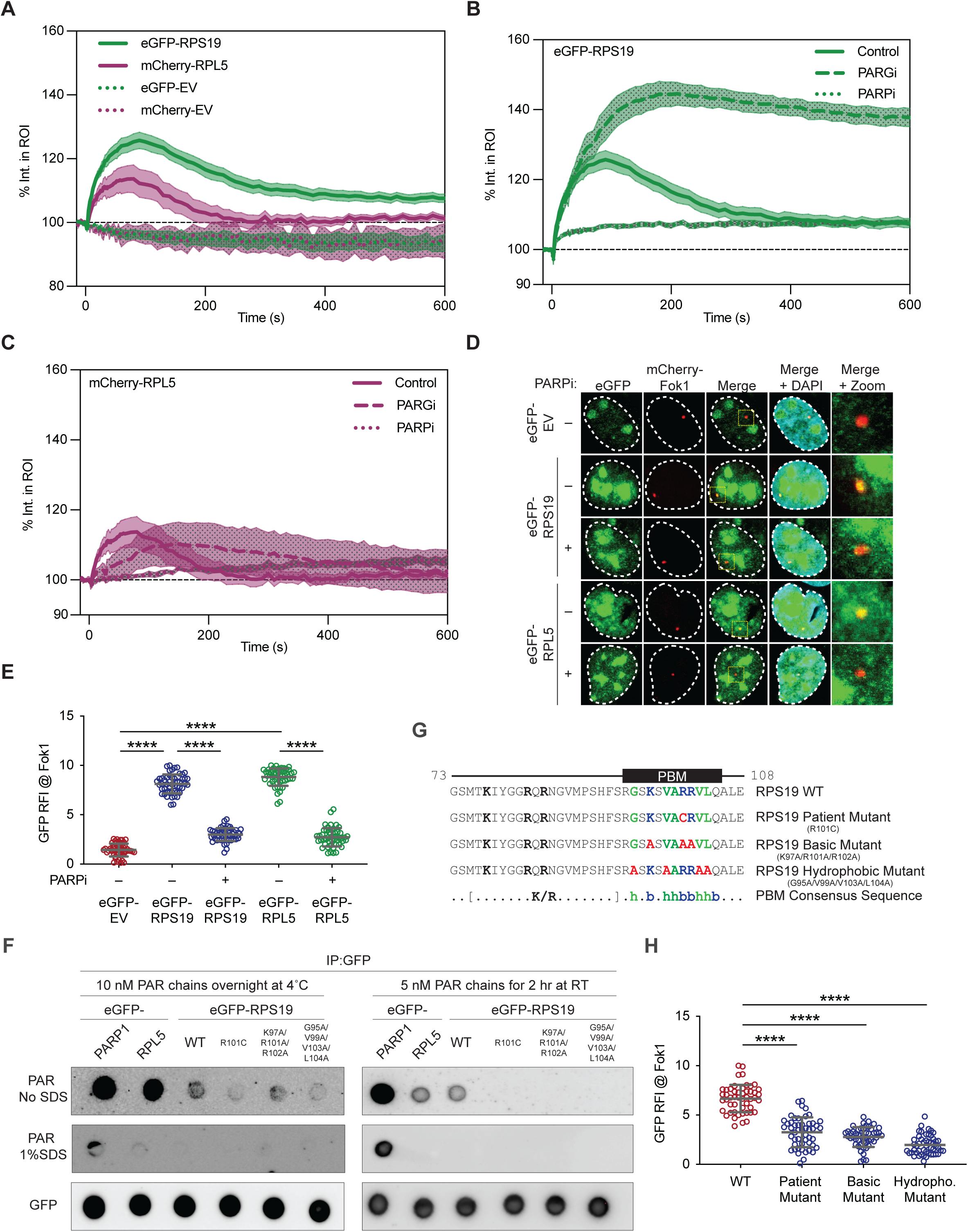
RPS19 and RPL5 are recruited to DSB sites in a PARP-dependent manner. (A) Quantification of percent fluorescent intensity within the region of interest (ROI) of U2OS cells transfected with eGFP-RPS19 (n = 8), mCherry-RPL5 (n = 8), or EVs (n = 3). (B) Quantification of percent fluorescent intensity within the ROI of eGFP-RPS19 transfected U2OS cells treated with PARP (n = 8) or PARG (n = 9) inhibitors. Control is the same as in A. (C) Quantification of percent fluorescent intensity within the ROI of mCherry-RPL5 transfected U2OS cells treated with PARP (n = 8) or PARG (n = 9) inhibitors. Control is the same as in A. (D) Representative images of eGFP-RPS19 and eGFP-RPL5 localization to mCherry-LacI-FokI DSB sites in U2OS cells with and without PARPi treatment. (E) Quantification of D. A minimum of 50 cells were analyzed per condition. (F) Blot of GFP-tagged proteins immunoprecipitated in 293T cells and incubated with 10 nM PAR chains overnight at 4°C (left) or 5 nM PAR chains for 2 hours (right) with or without 1% SDS and probed with a PAR antibody. GFP is used as loading control. (G) Schema of mutations generated within the potential PAR-binding motif of RPS19. (H) Quantification of eGFP-RPS19 WT and mutants localization to mCherry-LacI-FokI DSB sites in U2OS cells. A minimum of 49 cells were analyzed per condition. Data in A–C represent mean ± SEM. Data in E and H represent mean ± SD compared using unpaired t-tests. \*\*\*\**P* < 0.0001

Given that RPS19 and RPL5’s recruitment to DSBs depended on PARP activity, we sought to determine whether they bind poly(ADP) ribose (PAR) or are themselves PARylated. To test this, eGFP-RPS19 and eGFP-RPL5 immunoprecipitates (IPs) were applied to nitrocellulose membranes, which were then incubated with PAR chains under native (without SDS) or denaturing (+1% SDS) conditions, followed by immunoblotting. We detected binding to PAR chains by both RPS19 and RPL5 under native conditions, though to a greater extent by RPL5 (Figure 5F). Using more stringent incubation conditions, the binding to PAR was comparable. Importantly, the interaction was abolished under denaturing conditions regardless of the incubation conditions. These data indicate that RPS19 and RPL5 interact with PAR noncovalently.

Further supporting RPS19’s binding to PAR chains, we identified sequences that conform remarkably to the consensus PAR binding motif (PBM) for DDR proteins (Figure 5G).^35^ To investigate if the potential PBM in RPS19 is functional, we generated RPS19 mutants with alanine substitutions of the PBM basic or hydrophobic residues or bearing a mutation identified in a patient with DBA (p.R101C).^36^ We found that each of these mutants had reduced PAR binding (Figure 5F), confirming the functionality of the PBM. To assess if PAR binding is required for the recruitment of RPS19 to DNA breaks, we expressed eGFP-RPS19 WT and mutants in cells expressing a mCherry-LacI-FokI nuclease and found that only WT eGFP-RPS19 localized to FokI sites (Figure 5H; supplemental Figure 7B). Together, these data suggest that the noncovalent interaction of RPS19 with PAR chains via its PBM is necessary for its recruitment to DSBs.

We also investigated whether the recruitment of either eGFP-RPS19 or mCherry-RPL5 to sites of DSBs was dependent on other factors involved in the DDR. We found that their recruitment was unaffected by the ATR inhibitor VE-821 (supplemental Figure 7C–D). In addition, while eGFP-RPS19 and mCherry-RPL5 recruitment to sites of damage had similar kinetics between control and ATM inhibitor KU-60019 treated cells (supplemental Figure 7E–F), the percent fluorescence intensities for both proteins were reduced, suggesting an influence of ATM signaling on the recruitment of both RPS19 and RPL5 to DSBs.

### Recruitment of RPL5 is influenced by p53

As noted above, some ribosomal proteins are effectors of ribosomal stress through stabilizing p53.^37^ To investigate if the recruitment of RPS19 and RPL5 to sites of DSBs was influenced by p53, we first transfected Saos2 cells, which are p53-null, with eGFP-RPS19 and mCherry-RPL5 and monitored their recruitment to sites of laser microirradiation. We found rapid recruitment of RPS19 in Saos2 cells, like its recruitment in p53-positive U2OS cells (compare Figure 6A–B with Figure 5A; supplemental Figure 7A). In contrast, unlike its behavior in U2OS cells (Figure 5A; supplemental Figure 7A), we found that mCherry-RPL5 recruitment was undetectable in Saos2 cells (Figure 6A–C). To determine if this was due to an absence of p53 function, we co-expressed mCherry-RPL5 with a plasmid expressing WT p53, a plasmid expressing the p53 mutant p.R248Q, which results in loss of p53 binding to DNA,^38^ and a plasmid expressing a C-terminal domain truncation deletion mutant of p53 (ΔCTD), which eliminates its rapid association with DSBs but renders it transcriptionally hyperactive.^21,39,40^ While RPL5 recruitment in Saos2 cells remained undetectable when p53-R248Q was overexpressed, it was evident with p53 WT and ΔCTD overexpression (Figure 6D–E). To corroborate these findings and to overcome possible cell line differences between U2OS and Saos2 cells that extend beyond p53 status, we knocked down p53 in U2OS cells and monitored the recruitment of RPS19 and RPL5 to DSBs (Figure 6F–H). Similar to what was observed in Saos2 cells, RPS19 was rapidly recruited to DSBs, and the recruitment of RPL5 was significantly reduced in U2OS cells treated with a p53 siRNA compared to a control siRNA. These results confirm that p53 is required for the recruitment of RPL5 but not RPS19 to DSBs.

**Figure 6.**
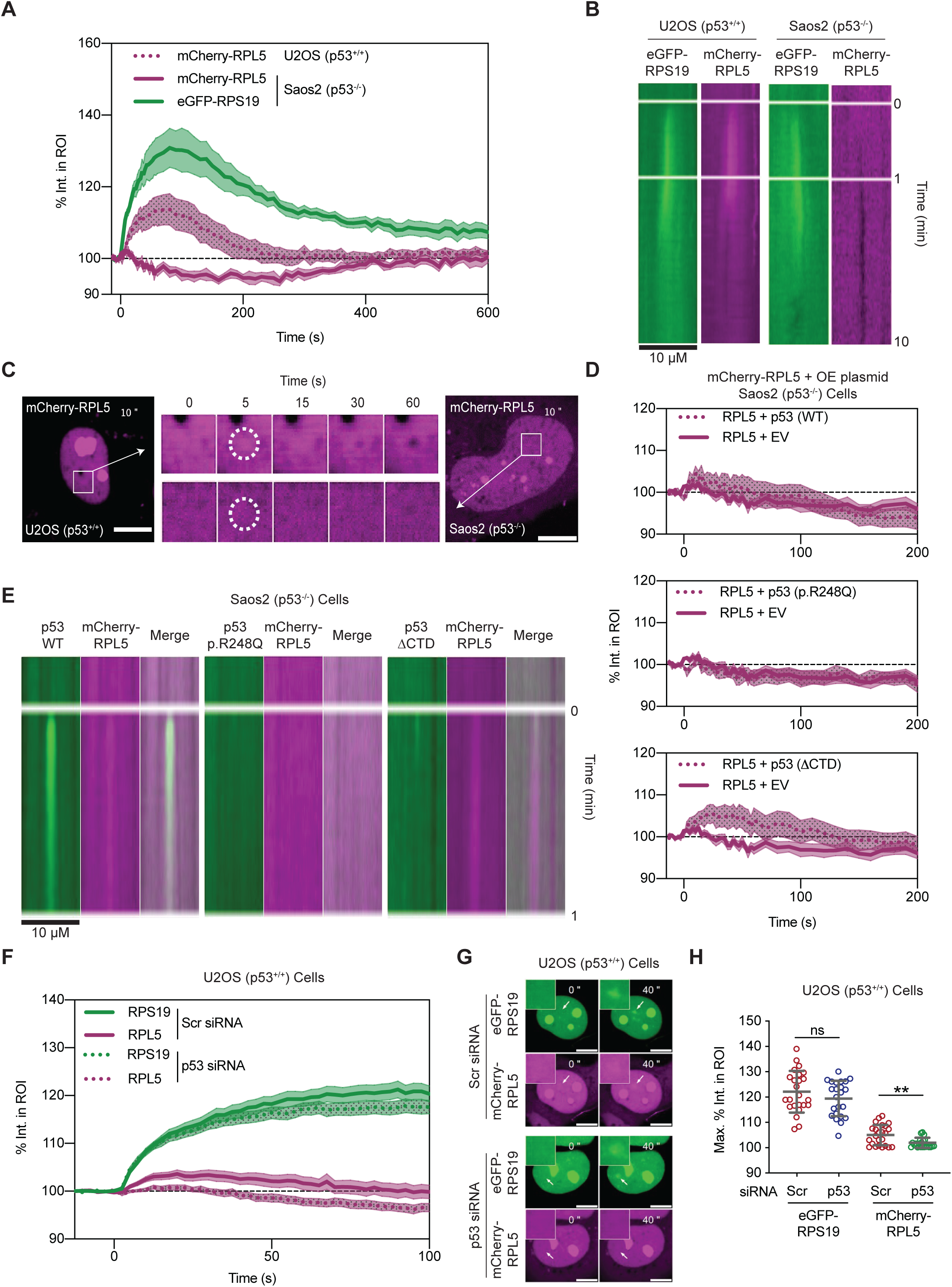
RPL5 but not RPS19 recruitment is dependent on p53. (A) Quantification of percent fluorescent intensity within the ROI of U2OS cells transfected with mCherry-RPL5 (n = 8) or Saos2 cells transfected mCherry-RPL5 (n = 7) or eGFP-RPS19 (n = 7). mCherry-RPL5 U2OS same as in Figure 6A. (B) Kymograph of eGFP-RPS19 and mCherry-RPL5 spatial distribution in U2OS and Saos2 cells over 1 min. (C) Representative montage showing mCherry-RPL5 recruitment in U2OS cells which is absent in Saos2 cells. White circle denotes the area of laser irradiation. (D) Quantification of percent fluorescent intensity within the ROI of Saos2 cells transfected with mCherry-RPL5 (n = 9) with WT p53 (n = 8), p53 p.R248Q (n = 7), or p53 ΔCTD (n = 9). (E) Kymograph of eGFP-p53 WT, eGFP-p53 p.R248Q, or eGFP-p53 ΔCTD and mCherry-RPL5 spatial distribution in Saos2 cells over 10 min. (F) Quantification of percent fluorescent intensity within the ROI of U2OS cells transfected with eGFP-RPS19 (n = 24) and Scr siRNA, mCherry-RPL5 (n = 24) and Scr siRNA, eGFP-RPS19 (n = 23) and p53 siRNA, mCherry-RPL5 (n = 23) and p53 siRNA. (G) Representative montage showing eGFP-RPS19 recruitment and lack of mCherry-RPL5 recruitment in U2OS cells treated with p53 siRNA. White arrow denotes the area of laser irradiation. (H) Quantification of the peak normalized intensity within the ROI for eGFP-RPS19 and mCherry-RPL5 recruitment with Scr or p53 siRNA treatment. Data in A, D, and F represent mean ± SEM. Data in H represent unpaired t-tests. ns not significant; \*\**P* < 0.01.

### RPL5 interacts with p53, and its recruitment to DSBs is disrupted when its binding to 5S rRNA is impaired

To strengthen the roles RPS19 and RPL5 have in DSB repair, we assessed if they interact with known repair factors. We found that both proteins interacted with the NHEJ factor Ku70 and histone H2A, which, when modified, can promote DSB signaling (Figure 7A; supplemental Figure 8A–B). Given p53’s influence on RPL5’s recruitment to DSBs, we also wondered if p53 and RPL5 interacted. We found that endogenous p53 was enriched in eGFP-RPL5 but not eGFP-RPS19 IPs relative to the eGFP-EV control IPs (Figure 7A; supplemental Figure 8A–B). However, transient transfection of eGFP-RPL5 led to a marked increase in p53, raising the possibility that the increased p53 protein level drove a nonspecific interaction. To address this, we treated transfected cells with 200 nM RG7388, which inhibits p53-MDM2 interaction and, thereby, stabilizes p53.^41^ We found that the resulting increase in p53 level with RG7388 treatment did not enhance the presence of p53 in the eGFP-EV IP, confirming that the interaction of p53 with RPL5 was specific (Figure 7B–C; supplemental Figure 8C). Notably, however, we also observed that RG7388 reduced the p53-RPL5 interaction suggesting that MDM2 influenced the association. Consistent with this, we found that the RPL5 mutant (p.N94D), which is impaired for 5S rRNA binding and MDM2 association,^42^ did not co-localize with FokI-induced DSBs (Figure 7D–E). We also found that RPL5-KD cells had decreased p53 protein levels as previously reported (Figure 7F–G);^43^ however, p53-KD increased both HR and total-EJ repair pathways (supplemental Figure 9). These results, combined with the impact of p53 on RPL5’s recruitment to DSBs (Figure 6), suggest an interplay between RPL5 and p53 that serves to suppress EJ repair.

**Figure 7.**
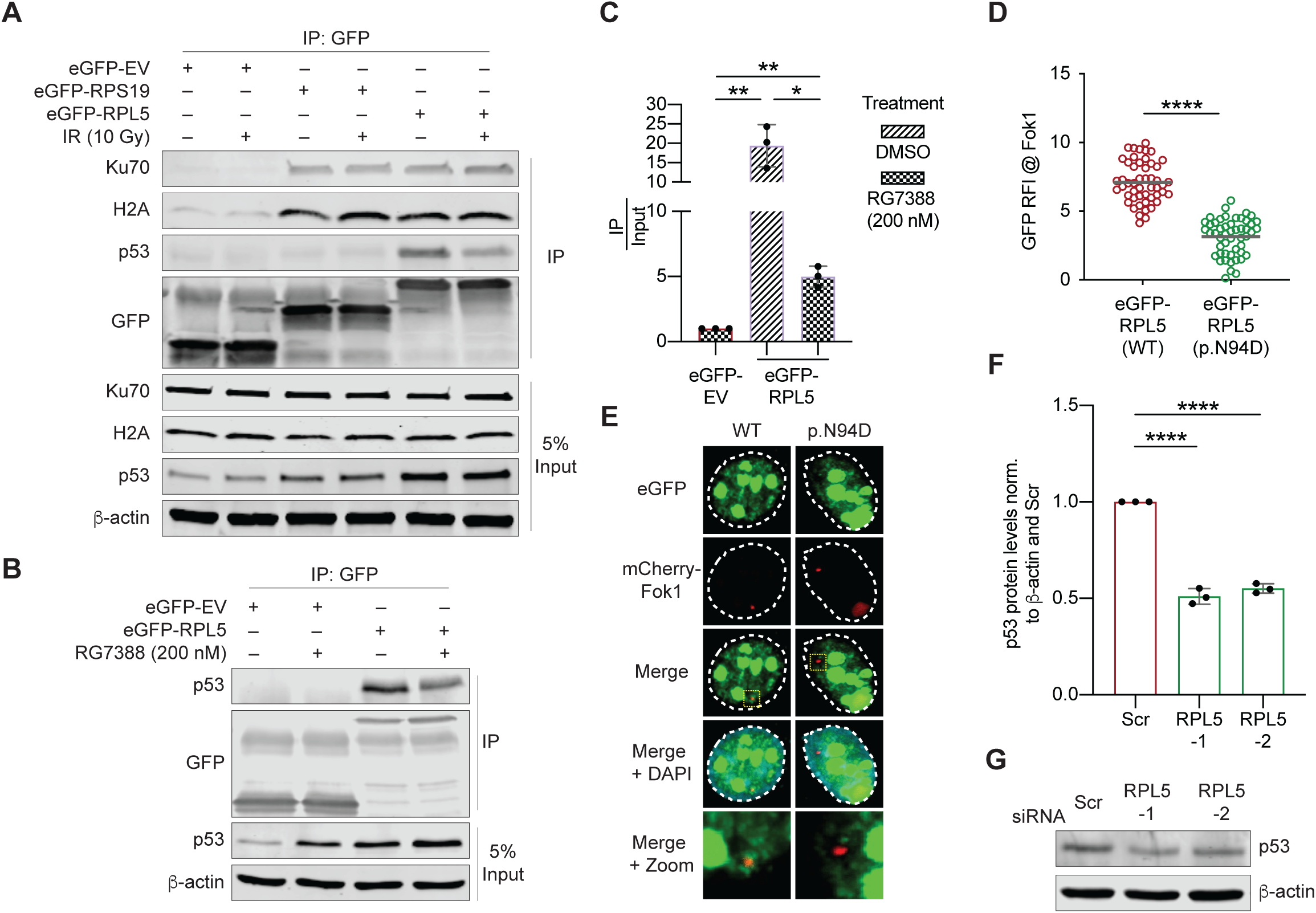
RPL5 interacts p53 in a MDM2-dependent manner and its association with DSBs involves the 5S RNP and possibly MDM2. (A) GFP immunoprecipitation (IP) of eGFP-RPS19 and eGFP-RPL5 transfected U2OS cells treated with or without 10 Gy IR. GFP is used as an IP control. β-actin is used as a loading control. (B) GFP IP of eGFP-RPL5 transfected U2OS cells treated with or without 200 nM of the MDM2 inhibitor RG7388. GFP is used as an IP control. β-actin is used as a loading control. (C) Quantification of B relative to input. Data represent three biological replicates. (D) Quantification of eGFP-RPL5 WT and p.N94D mutant localization to mCherry-LacI-FokI DSB sites in U2OS cells. Fifty cells were analyzed per condition. (E) Representative images of blots quantified in D. (F) Quantification of p53 protein levels in RPL5-KD U2OS cells normalized to β-actin and Scr. Data represent three biological replicates (G) Representative western blot of data shown in F. β-actin used as a loading control, Data in C–D represent mean ± SD compared using unpaired t-tests. Data in F represent mean ± SD compared using 1-way ANOVA with Dunnett’s multiple comparisons test. \**P* < 0.05; \*\**P* < 0.01; \*\*\*\**P* < 0.0001.

## DISCUSSION

Here, we identify RPS19 and RPL5 in supporting DNA DSB repair, providing, for the first time, evidence for DSB repair defects associated with deficiencies of these major DBA-associated RPs. Despite both being core ribosome constituents and associated with similar impacts on the proteome when depleted,^4^ we found their depletion resulted in different effects on the repair of DSBs, with RPS19-KD impairing pathways requiring extensive resection (HR and SSA) and RPL5-KD increasing repair via end-joining. HR deficiency (HRD) has been associated with various cancers and germline cancer predisposition syndromes, and many groups have identified agents that leverage HRD to treat cancer.^44,45^ Alt-EJ is associated with genome instability as it results in base pair deletions as well as chromosomal rearrangements and translocations.^46^ Additionally, loss of tumor suppressors such as TGFβ results in increased alt-EJ, which may promote cancer development.^47^ Our data suggest that a DSB repair pathway utilization shift, with reduced high-fidelity HR or more mutagenic end-joining, could increase mutation accumulation in cells with RPS19- or RPL5-deficiency, respectively. Thus, we propose that disruption of DSB repair may underlie, at least in part, the increased risk of cancer in individuals with DBA due to *RPS19* or *RPL5* germline pathogenic variants and, as we discuss below, potentially due to additional RP genes.

How might RPS19 and RPL5 impact DSB repair? Of the 22 proteins involved in the DDR or DSB repair pathways that we examined, the levels of only a few, though each notable, were significantly impacted by RPS19- or RPL5-KD. RAD51 levels were decreased and LIG4/XRCC4 levels were increased by both RPS19- and RPL5-KD, and BRCA2 and PARP1 were lowered by RPS19-KD but inconsistently by RPL5-KD. While RPS19 deficiency has been shown to decrease the TE of specific transcripts,^4,48^ we found the TE of only RAD51 mRNA had decreased, and this was accompanied by an increase in the level of RAD51 mRNA and an increase in RAD51 proteasomal degradation. Moreover, we observed an increase in PARP1 and no change in BRCA2 TE in RPS19-KD cells but lower levels of both proteins. Altogether, these data highlight that processes other than or in addition to translation may be altered with RP deficiencies to affect the level of proteins that lead to cellular phenotypes associated with DBA. Irrespective of the mechanism(s) altering protein levels, the combined reduction in PARP1, BRCA2, and RAD51 alone could underlie the reduced HR efficiency in RPS19-KD cells. Not analyzed in our study, reduction in other factors could also play a role. Indeed, we observed that RAD51 levels were also reduced in RPL5-KD cells, yet HR remained unchanged.

RPS19 and RPL5 were rapidly recruited to DSBs, suggesting that they might impact repair, at least in part, by direct interactions at these sites. Like their effects on DSB repair pathways, we discerned differences in features of their recruitment, such as the sensitivity to PARG inhibition. RPS19 contained a PBM characteristic of DDR factors, and this motif was required for PAR chain binding and recruitment to DSBs. While we have not identified the specific PARylated factor(s) to which it binds, this finding is consistent with previous studies showing that many known DSB repair factors are recruited to break sites via interactions with PARylated proteins, including histones,^49^ further substantiating a role in DSB repair.

RPS19 and RPL5’s recruitment to DSBs were also distinguishable by the critical influence of p53 on RPL5’s recruitment. RPL5-KD resulted in a decrease in p53 protein levels, and p53-KD resulted in increased total-EJ and HR. Though RPL5-KD increased total-EJ but not HR, HR was increased when LIG4/XRCC4 were restored to normal levels. Thus, RPL5 and p53 mirror each other with regard to suppressing these pathways, corresponding to the dependence of RPL5 on p53 for its recruitment to DSBs. We also found RPL5, but not RPS19, in association with p53. While the binding of MDM2 to 5S RNP inhibits ubiquitination of p53 and p53 stabilization, we found that inhibiting MDM2’s association with p53 via RG7388 also reduced RPL5’s association with p53. Additionally, we observed that the RPL5 N94D mutation, which impairs MDM2’s association with 5S RNP, also impairs RPL5’s association with DSBs. Together, these results suggest that the interplay between RPL5-MDM2-p53 not only regulates p53 levels but also has a direct influence on RPL5’s recruitment and role in suppressing DSB repair.

Ribosomal protein RPL6, which has not been implicated in DBA, is also recruited to DSBs. Once there, it promotes the binding of MDC1 to γ-H2AX, leading to the eventual ubiquitination of γ-H2AX and recruitment of DNA repair proteins, such as 53BP1.^15^ While we have not determined the impacts of RPS19- or RPL5-KD on recruitment of these factors, we have found the effects of each on DSB repair pathways are distinct from that of RPL6-KD in which both HR and NHEJ are reduced. RPS3, a non-DBA-associated RP, has been shown to interact with Ku heterodimers and slow the ligation of break ends in NHEJ.^50^ Like RPS3, RPL5 may directly inhibit end-joining repair at DSBs as RPL5-KD enhances the repair efficiency of total-EJ and alt-EJ and eGFP-RPL5 interacts with Ku70; however, the interaction of Ku70 and RPL5 most likely does not contribute to the impact RPL5-KD has on end-joining repair as RPS19 also interacts with Ku70. Overall, further investigations are needed to elucidate how RPS19 and RPL5’s recruitment to DSBs impacts different DSB repair pathways.

Warner and McIntosh proposed three criteria for an RP having an extraribosomal capacity: demonstrations that the RP interacts with a nonribosomal factor, that this occurs while away from the ribosome, and that this has a physiological effect on the cell.^51,52^ In support of extraribosomal functions, we have discovered that RPS19 and RPL5 localize at sites of DSBs and interact with known DSB repair factors. In addition, differences in their recruitment in Saos2 cells, as well as dependencies on PARP and p53, argue that their recruitment is independent of the ribosome. Although we have demonstrated differential effects of RPS19- and RPL5-KD on DSB repair pathways, connecting these effects to their recruitment to sites of DSBs will be required to definitively attribute an extraribosomal function in DSB repair for these proteins. Whether extraribosomal or not, the haploinsufficiency of these proteins may challenge the repair of the estimated 50 DSBs that occur per cell cycle,^53^ ultimately giving rise to cancer-driving mutations. These findings provide new insights and a model to test and understand the mechanisms that drive cancer predisposition in DBA.

## ACKNOWLEDGMENTS

We thank the research subjects for their participation and the clinical research staff for their assistance. We thank the members of the Bertuch lab for helpful discussions throughout this project and critical reading of the manuscript, Dr. Amos Gaikwad and Ms. Tatiana Goltsova at the Texas Children’s Hospital Flow Cytometry Core for their assistance with the flow cytometry experiments, and Dr. Fabio Stossi, Ms. Hannah Johnson, and Ms. Elina Mosa at Baylor College of Medicine’s (BCM) Integrated Microscopy Core for their assistance with the immunofluorescence experiments. We also thank Dr. Eveline Barbieri for the MDM2 inhibitor and Amber Wolf for her guidance with using the BD Symphony A1 flow cytometer. We thank Dr. Jeffrey M. Rosen (J.M.R.) for assistance with the polysome profiling experiments. This work was supported by the US Department of Defense (W81XWH-21-1-0415 to A.A.B.) and the National Institutes of Health (T32DK060445 and T32GM136554 to N.F.D.; R35GM150992 to N.C.; R01CA198279 and R01CA250905 to K.M.M.; R01GM149683 to Y.H.W.; DK56338, CA125123, and ES030285 to BCM’s Integrated Microscopy Core; CA125123 and RR024574 to BCM’s Cytometry and Cell Sorting Core), and the Cancer Prevention Research Institute of Texas (RP220330 to K.M.M.; RR180025 to Y.H.W.; RP220468 to J.M.R.; RP150578, and RP170719 to BCM’s Integrated Microscopy Core; RP180672 to BCM’s Cytometry and Cell Sorting Core), the Breast Cancer SPORE Career Enhancement Award (to N.Z.), and the American Cancer Society (PF-22-092-01-DMC to A.S.) The content is solely the responsibility of the authors and does not necessarily represent the official views of the US Department of Defense, the National Institutes of Health, the Cancer Prevention Research Institute of Texas, or the American Cancer Society. N.F.D. is a Ph.D. candidate at Baylor College of Medicine. This work is submitted in partial fulfillment of the requirement for the Ph.D.

## AUTHORSHIP

Contribution: N.F.D., E.A., and A.A.B. conceptualization; N.F.D., E.A., A.S., C.L.W., E.B.K., N.Z., N.C., and Y.H.W. investigation; N.F.D. writing – original draft; E.A., A.S., C.L.W., E.B.K., N.Z., N.C., K.M.M., Y.H.W., and A.A.B. writing – review and editing; K.M.M. and A.A.B. supervision; A.A.B. funding acquisition.

Conflict-of-interest disclosure: The authors declare no competing financial interests.

Correspondence: Alison A. Bertuch, M.D., Ph.D., Texas Children’s Hospital, 1102 Bates, Ste 1200, Houston, TX 77030.

