## Supplemental Materials, Methods, and Figures for "*RPS19* and *RPL5*, the most commonly mutated genes in Diamond Blackfan anemia, impact DNA double-strand break repair"

#### SUPPLEMENTAL METHODS

##### Cell lines and culture and plasmids

Lymphoblastoid cell lines (LCLs) were generated under Baylor College of Medicine (BCM) IRB-approved study H-17698 using the tissue culture core laboratory within the Department of Molecular and Human Genetics, BCM. LCLs were cultured in RPMI 1640 medium containing L-glutamine and 10% FBS in 5% $CO_2$. U2OS cells were maintained in McCoy's medium supplemented with 10% FBS in 5% $CO_2$. CD34+ cells were isolated from healthy donor bone marrow aspirate using Dynabeads CD34 (Invitrogen, 11301D) and grown in StemSpan SFEM (StemCell, 09650) supplemented with human SCF, human FLT-3, human TPO, and human IL3 (PeproTech).

DR-GFP-U2OS reporter cells were generated by electroporating (Lonza, VCA-1003) U2OS cells with Sac1/Kpn1 linearized pHPRT-DRGFP (Addgene #26476) DNA.[1] Puromycin-resistant clones were pooled 10–15 days later and used for HR assays. EJ5-GFP-U2OS, SA-GFP-U2OS, and EJ2-GFP-U2OS reporter cells were similarly generated using XhoI linearized pimEJ5-GFP (Addgene #44026), Hpa1 linearized hprtSA-GFP (Addgene #41594), and Sac1/Kpn1 linearized EJ2-GFP (Addgene #44025) DNA, respectively.[2,3] DR-GFP-HCT116 and EJ5-GFP-HCT116 were generated similarly as described above.

The *RPS19* siRNA-resistant plasmid (pAB1180) was generated by cloning a gBlock (Integrated DNA Technologies) into an *RPS19-3xFLAG* plasmid (pAB1162). pAB1162 was created via Gateway Cloning (entry clone: *RPS19* in pDONR223 [AP404 from BCM's Cell-Based Assay Screening Service Core], destination vector: pGCS-C3m(3xFLAG) [Addgene #85726]). The *RPS19* gBlock containing silent mutations in the region complementary to the *RPS19* siRNAs was digested with SacI/NotI and subcloned pAB1162 to replace the wild-type sequence. The *6xMYC-LIG4* plasmid (pAB1313) was generated by subcloning PCR-amplified *LIG4* cDNA

(Addgene #37150) into pCS2+MT (AP106). The *2xFLAG-XRCC4* plasmid (pAB1312) was generated by subcloning PCR-amplified XRCC4 cDNA (Addgene #46959) into pcDNA3.1 (AP2). The *eGFP-RPS19* plasmid (pAB1222) was generated by subcloning PCR-amplified *RPS19* cDNA (AP404) into pEGFP-C1 (AP134). The *eGFP-RPL5* plasmid (pAB1249) was generated by subcloning an XhoI/BamHI digested gBlock (Integrated DNA Technologies) containing *RPL5* cDNA into p*EGFP*-C1 (AP134). The *mCherry-RPL5* plasmid (pAB1262) was generated by subcloning a pCAGGS-mCherry fragment PCR-amplified from Addgene plasmid #41583 into *eGFP-RPL5* (pAB1249). The *eGFP-RPS19* patient mutant (p.R101C [pAB1284]) was generated by subcloning an HindIII/NotI digested gBlock (Integrated DNA Technologies) containing *RPS19* (c.301C>T) cDNA into *eGFP-RPS19* (pAB1222) to replace the wild-type sequence. The *eGFP-RPS19* basic mutant (p.K97A, p.R101A, p.R102A [pAB1285]) was generated by subcloning an HindIII/NotI digested gBlock (Integrated DNA Technologies) containing *RPS19* (c.288A>G, c.289A>C, c.290G>T, c.300C>A, c.301G>A, c.302C>G, c.303C>A, c.304G>A) cDNA into eGFP-RPS19 (pAB1222) to replace the wild-type sequence. The *eGFP-RPS19* hydrophobic mutant (p.G95A, p.V99A, p.V103A, p.L104A [pAB1268]) was generated by subcloning an XmnI/BamHI digested gBlock (Integrated DNA Technologies) containing *RPS19* (c.284G>C, c.296T>C, c.296T>C, c.297G>A, c.310C>G, c.311T>C) cDNA into eGFP-RPS19 (pAB1222) to replace the wild-type sequence. The *eGFP-PARP* plasmid was generated by gateway cloning PARP1 into the pcDNA6.2 N-EmGFP-DEST backbone. The *eGFP-RPL5* MDM2 binding mutant (p.N94D [pAB1270]) was generated by subcloning an EcoRV/BamHI digested gBlock (Integrated DNA Technologies) containing *RPL5* (c.280A>G) cDNA into *eGFP-RPL5* (pAB1249) to replace the wild-type sequence. Plasmids encoding p53-WT-EGFP and mutant derivatives were generated as previously described.[4]

**Immunofluorescence (IF) assays**

γ-H2AX foci: Lymphoblastoid cells were grown to >75% confluence on 6-well dishes and treated with a single dose of 2 Gy ionizing radiation (IR). Cells were harvested, fixed in 4% paraformaldehyde, quenched with $NH_4Cl$ 0.1 M in phosphate-buffered saline (PBS), and spotted in 384-well glass bottom plates by cytospin at 1000 rpm for 3 minutes. Cells were permeabilized with 0.1% Triton-X100. After blocking in 5% milk in Tris-buffered saline + 1% tween (TBS-T), cells were incubated with anti-phospho-histone H2A.X (Ser139; γ-H2AX) antibody (Millipore, clone JBW301) at 1:1000 dilution overnight. After washing, cells were incubated at room temperature (RT) with Alexa Fluor 488-conjugated IgG antibody (Life Technologies, A28175) at 1:1000 dilution for 30 minutes and counterstained with DAPI (2 μg/mL). The number of γ-H2AX foci per nucleus was measured from cells imaged with a GE Healthcare DeltaVision deconvolution microscope using an Olympus 40×/0.95NA PlanApo objective. Z-stacks (0.35 μm steps) were acquired, covering the entire nucleus, and projected by maximum intensity after restorative deconvolution. Foci were identified using a local maxima detection algorithm after denoising and Gaussian filtering the γ-H2AX. This analysis was performed with MATLAB version 2016b.

Nuclear RPA2 integrated intensity: U2OS cells were plated in 6-well plates. Twenty-four hours later 25 nM RPS19 or scrambled (Scr) siRNA (see supplemental Table 2 for siRNA sequence) was transfected using RNAi Max (Invitrogen, 13778). Forty-eight hours later, cells were plated on coverslips. The following day, cells were treated with 10 Gy of IR or reserved for untreated (UT) controls. Cells were harvested, fixed in 4% paraformaldehyde, permeabilized in 0.2% Triton X-100, and blocked in 3% BSA. Cells were incubated overnight in anti-RPA2 antibody (Invitrogen, PA5-22256) at 1:500 dilution. The following day, cells were incubated in Alexa Flour 488-conjugated antibody (Invitrogen, A11034) at 1:1000 dilution and counterstained with 0.1 μg/mL Hoechst 33258. Cells were mounted in SlowFade Diamond (Thermo, S36972). The RPA2 intensity was measured using a GE Healthcare DeltaVision deconvolution microscope with an

Olympus 20× (0.75NA) U-Plan S-Apo objective. Z-stacks (0.70 μm steps) were acquired, covering the entire nucleus, and projected by maximum intensity after restorative deconvolution. Nuclear masks were defined based on the DAPI signal, and RPA2 nuclear intensity was assessed using CellProfiler version 4.2.1.

RAD51 foci: RAD51 IF was conducted similarly to the RPA2 IF, except an anti-RAD51 primary antibody (Abcam, ab176458) at 1:500 dilution was used. The RAD51 foci were measured using an Olympus IX83 inverted microscope with an Olympus 60× (1.42NA) PLAPON oil objective. Z-stacks (0.24 μm steps) were acquired, covering the entire nucleus, and projected by maximum intensity after restorative deconvolution. Nuclear masks were defined based on the DAPI signal, and RAD51 foci were counted using CellProfiler version 4.2.1.

**Neutral comet assays**

The Bio-Techne manufacturer’s protocol (4250-050-5) was used. RPS19-1, RPS19-2, RPL5-1, RPL5-2, or Scr siRNA (25 nM) was electroporated into CD34+ cells. Following 48 hours incubation in StemSpan SFEM (StemCell, 09650) supplemented with human SCF, human FLT-3, human TPO, and human IL3 (PeproTech), cells were treated with 10 Gy IR using a Rad Source Pro Biological Irradiator Model 2000 (X-ray, 25 mA, 160 kV). The cell solutions were added to a slide coated with 0.65% agarose. Following lysis, slides were ran in the electrophoresis chamber filled with Neutral Electrophoresis Buffer (see manufacturer’s protocol). DNA was precipitated and staining solution was added. Slides were imaged immediately after using an Olympus IX71 fluorescence microscope with an Olympus 20× (0.45NA) LUCPlanFL N objective. Images were analyzed using ImageJ with the OpenComet software v1.3.1.

**Immunoblotting**

Cells were lysed in RIPA buffer (50 mM Tris-HCl pH 8.0, 150 mM NaCl, 1% NP40 v/v, 0.5% Na deoxycholate, 1% SDS, and 5 mM EDTA) with 1x protease inhibitor cocktail III (Calbiochem, 539134) and 1x PMSF (Sigma, 93482) for 10 minutes on ice, sonicated for 5 minutes (30 seconds on, 30 seconds off) using a Bioruptor UCD-200 sonicator, and centrifuged at 20,000 rcf for 20 minutes at 4˚C. Protein concentration was determined using the bicinchoninic acid (BCA) protein assay kit (Pierce, 23225). Depending on the amount of protein isolated across the sample set, an equivalent amount of protein, ranging from 20 to 50 µg, was loaded per sample in 1x protein dye. The samples were run on 4-20% Mini-Protean TGX precast gels (BioRad, 4561094) and transferred to Immobilon-FL PVDF membranes (Millipore, IPFL00010) and probed with antibodies as indicated (supplemental Table 3) in Intercept Blocking Buffer (LI-COR, 927-70001) overnight at 4˚C. Membranes were washed in PBS + 0.1% Tween (PBS-T) and detected with secondaries as indicated in Intercept Blocking Buffer for 1 hour at RT. Membranes were washed in PBS-T. Protein band quantification was performed using a LI-COR Odyssey CLx Infrared Imaging System and Image Studio Lite software.

For BRCA2, PARP1, and PARP2: Cell pellets were suspended in PBS. An equal volume of 2X lysis buffer (100 mM Tris pH 7.0, 4% SDS, and 12% β-mercaptoethanol) was added. Samples were boiled at >95˚C for 10 minutes and centrifuged at 12,000 rcf for 10 minutes. Protein concentration was measured using a nanodrop. Samples were run on a freshly made 4-18% gradient gel and transferred to a 0.45 PVDF membrane (Millipore, IPVH00010) using the BioRad Trans-Blot Turbo system (BioRad, 1704150) according to the manufactures instructions. Membranes were blocked with 3%BSA in TBS-T for 10 minutes and then blotted overnight at 4˚C with the indicated antibodies (supplemental Table 3) diluted in 5%BSA/TBS-T. Membranes were washed in TBS-T and detected with secondaries (diluted in 3%BSA/TBS-T) for 1 hour at RT. Signals were detected with Amersham ECL Prime Western Blotting Detection Reagent (Cytiva,

RPN2232) and visualized on a BioRad ChemiDoc MP. Protein band quantification was performed using ImageJ with β-tubulin used as a loading control.

**HR, total-EJ, SSA, and alt-EJ reporter assays**

Reporter cells were plated in 6-well plates and transfected with 5 µL Lipofectamine RNAi MAX transfection reagent (Invitrogen, 13778) and siRNAs (Invitrogen) at 25 nM final concentrations according to the manufacturer's instructions. Forty-eight hours later, the cells were co-electroporated (Lonza, VCA-1003) with 10 µg pCBA-SceI (Addgene, plasmid #26477) and 1 µg pCAGGS-mCherry (Addgene, plasmid #41583) expression plasmids. The cells were harvested two days later and subjected to flow cytometry analysis to detect GFP-positive and mCherry-positive cells using a BD LSRII flow cytometer and BD FlowJo software. The repair efficiency was scored as the percentage of GFP- to mCherry-positive cells.

For LIG4-KD experiments: Twenty-four hours-post knockdown as described above, cells were transfected with a LIG4 siRNA. Twenty-four hours later, cells were co-electroporated with I-SceI and mCherry expression plasmids, and the cells were harvested two days later and subjected to flow cytometry analysis as described above.

For LIG4 and XRCC4 overexpression experiments: *6xMYC-LIG4* and *2xFLAG-XRCC4* (2 µg each) expression plasmids were co-transfected using Lipofectamine® (Invitrogen, 18324012) and PLUS (Invitrogen, 11514015) reagents into HR and total-EJ reporter cells. Forty-eight hours later, cells were co-electroporated with I-SceI and mCherry, and the cells were harvested two days later and subjected to flow cytometry analysis as described above. Samples were analyzed using the BD FACS Symphony A1 flow cytometer due to retirement of the BD LSRII flow cytometer.

**Cell cycle analysis**

Seventy-two hours following siRNA transfections as described above, cells were grown for 2 hours in medium supplemented with 20 µM BrdU (Sigma, B5002). Cells were fixed with 70% ethanol, incubated at -20˚C for at least 16 hours, and then pelleted (here and throughout) by centrifugation at 600 rcf at RT. The DNA was denatured in 2 M HCl for 20 minutes at RT. Cells were pelleted, washed with PBS + 0.5% BSA, pelleted, and then incubated in 0.1 M sodium tetraborate (pH 8.5) for 2 minutes at RT. Cells were pelleted, washed, and pelleted as described above and then resuspended in 5 µL of FITC-labeled anti-BrdU antibody (BioLegend, 364104) for 1 hour at RT protected from light. Cells were pelleted, washed, and pelleted as described above and then resuspended in 50 µg/mL propidium iodide (BioLegend, 421301) for 1 hour at RT protected from light. Cells were pelleted and resuspended in 1x PBS and analyzed using a BD LSRII flow cytometer and BD FlowJo software.

**Cell proliferation**

Twenty-four hours following siRNA transfections as described above, HCT116 cells were plated in 96-well plates (1,000 cells/well) in technical triplicates. The plate was incubated at RT for 10 minutes and subsequently placed in 37˚C incubator overnight to ensure cell attachment. The next day, the plate was placed in the Incucyte S3 Instrument. Whole wells were imaged every 2 hours for 72 hours. The confluence of each well was assessed utilizing Incucyte v2022B Rev2 software, and the average confluence was calculated between the technical triplicates. The experiment was repeated two additional times.

**Polysome profiling**

Sucrose gradients (10–50%) were prepared in polysome buffer (20 mM Tris pH 7.4, 150 mM NaCl, 5 mM $MgCl2$) supplemented with 100 µg/mL cycloheximide (Sigma, C4859), 20 U/mL SUPERase•In RNase Inhibitor (Invitrogen, AM2696) and RNase-free sucrose (Sigma-Aldrich,

84097), and poured into polypropylene tubes (Beckman Coulter, 331374). Tubes were stored at 4˚C overnight.

Seventy-two hours following siRNA transfections as described above, cells were treated with 100 µg/mL cycloheximide (CHX) for 5 minutes, washed 1x in ice cold PBS containing 100 µg/mL cycloheximide, trypsinized with trypsin containing 100 µg/mL cycloheximide, and collected in media containing 100 µg/mL cycloheximide to stop trypsin activation. Cells were centrifuged and washed 1x in ice cold PBS containing 100 µg/mL cycloheximide. Cells were lysed in 500 µL polysome buffer supplemented 100 µg/mL cycloheximide, 1% Triton X-100 (Teknova, T1105), and 25 U/mL Turbo DNase I (Invitrogen, AM2238). Cells were then passed 5 times through a 20-gauge needle, 10 times through a 25-gauge needle, and 15 times through a 27-gauge needle and incubated on ice for 10 minutes. Cells were then centrifuged for 10 minutes at 14,000g at 4°C and layered on top of sucrose gradients.

The gradients were centrifuged at 35,000 rpm for 2 hours at 4°C using an SW41 Ti rotor (Beckman Coulter). The gradients were displaced into a UA-6 continuous UV detector (Teledyne ISCO) using a syringe pump (Brandel) containing Fluorinert FC-40 (Sigma-Aldrich, F9755) at a speed of 0.75 mL/min. The absorbance was recorded at an OD of 260 nm using the Logger Lite (v1.9.4) software (Vernier) and a total of 24 fractions with approximate 500 µL volume for each fraction were collected using the Foxy Jr fractionator.

Following fraction collection, 10% SDS was added to each fraction for a final concentration of 1%. The tubes were mixed and 750 µL of Trizol LS (Thermo, 10296028) were added to each fraction. As an RNA control, 500 pg of luciferase RNA spike-in (Promega, L4561) was added to each fraction. The RNA was extracted from each fraction and 6 µL of RNA was converted to cDNA using the High-Capacity cDNA Reverse Transcription Kit (Thermo, 4368814). The cDNA was diluted 10x before PCR analysis. Quantitative PCR was performed using primers listed in supplemental Table 4 and on a QuantFlex 6 system (Applied Biosystems) using PowerUp SYBR Green Master Mix (Applied Biosystems, A25742) according to the manufacturer's protocol. The relative changes in RNA abundance were calculated as described previously.[5]

**RNA analysis**

Seventy-two hours following siRNA transfections as described above, RNA was isolated from U2OS cells using the RNeasy® kit (QIAGEN, 74004) and cDNA synthesized using the qScript Flex cDNA Synthesis Kit using random primers (Quanta Biosciences, 95049). Quantitative PCR was performed using primers listed in supplemental Table 4 and on a QuantFlex 6 system (Applied Biosystems) using PowerUp SYBR Green Master Mix (Applied Biosystems, A25742) according to the manufacturer's protocol. The relative changes in RNA abundance were calculated by comparative $\Delta$Ct method with normalization to GAPDH.

**Laser microirradiation**

The laser microirradiation experiments were performed with a UVA laser (UGA-42 Caliburn, Rapp OptoElectronics: 355 nm; 1 kHz repetition rate; 4.2 ns pulse width) installed on an Olympus SpinSR-10 Yokogawa spinning disk confocal and captured by an ORCA Fusion sCMOS camera (Hamamatsu). The 355 nm laser beam was introduced to the sample through an Olympus 100× (1.5NA) UIS2 XLine Plan Apochromat (UPLAPO) oil objective with an ND2 filter. *eGFP-RPS19* and *mCherry-RPL5* plasmids were transiently transfected into U2OS cells using Lipofectamine 2000/3000. Cells were sensitized with 10 µg/mL Hoechst 33342 (Sigma-Aldrich) for 10 minutes within 12-18 hours post transfection. Localized DNA damage was introduced by exposing regions of interest (ROIs) within nuclei to the UVA laser for 250 ms at 0.2 nW laser power measured at the back focal plane of the objective. Image acquisition was taken in two stages. The first set of time-lapse images was taken at 2.5-second intervals for the first 1 minute following laser microirradiation to capture the rapid recruitment of RPS19 and RPL5. The rest of the movie was taken at 30-second intervals for another 9 minutes to monitor the dissipation of their recruitment.

For knockdown experiments followed by laser microirradiation, cells were transfected with Lipofectamine RNAi MAX for 24 hours before the second round of transfection with the plasmids of interest (RPS19 or RPL5) using Lipofectamine 2000. Cell culture media were replaced 6 hours post each transfection and the laser microirradiation was performed 48 hours after siRNA transfection as described above.

For inhibitor experiments followed by laser microirradiation, drug-containing media was added 6 hours after cells were transfected with indicated plasmids by Lipofectamine 2000, and the cells were imaged 12-24 hours later as described above.

**Fok1 co-localization assays**

U2OS reporter cells[6] were transfected with the indicated plasmids using Lipofectamine 3000 (Invitrogen, L3000008) according to the manufacturer's protocol. Where indicated, cells were treated for 12 hours with 10 μM olaparib (Cayman Chemical, 10621). Site-specific DSBs were induced by adding 1 μM Shield-1 (Takara, 632189) and 1 μM 4-hydroxytamoxifen (Sigma, H7904-5MG) for 5 hours before fixation.

Cells were fixed as previously described.[7] In brief, cells were pre-extracted with CSK buffer (10 mM PIPES, pH 6.8; 100 mM NaCl; 300 mM sucrose; 3 mM MgCl2; 1 mM EGTA; 0.5% (v/v) Triton X-100) for 5 minutes on ice, washed with PBS, and fixed with 2% (v/v) formalin for 15 minutes at RT. Coverslips were mounted onto slides with VECTASHIELD mounting medium with DAPI (Vector Laboratories). The GFP and mCherry signals were directly visualized by a FluoView 3000 confocal microscope (Olympus) using Tru-Spectral high-efficiency gallium arsenide phosphide spectral detectors with a 60× (1.42NA) Plan Apochromat oil objective. Image acquisition was performed with FV-10 ASW3.1 software (Olympus).

Quantification of relative fluorescence intensity (RFI) was performed using ImageJ software. The integrated density of GFP at the DSB was measured and corrected to account for the background in the image. Values were normalized on a 0-10 scale, with the brightest measurement as 10.

**PAR binding assay**

PAR chains were synthesized and binding assay were performed as previously described.[8] Briefly, GFP-tagged proteins were transfected into 293T cells with FuGENE HD (Promega, E2311) and allowed to express for 48 hours. Cells were lysed by rotating at 4ºC in FA lysis buffer (50 mM HEPES-KOH pH7.6, 140 mM NaCl, 1 mM EDTA pH8, 1% Triton X-100, 0.1% sodium deoxycholate) for 1 hour, GFP-tagged proteins were enriched by incubating lysates overnight at 4ºC with GFP-trap magnetic agarose (Chromotek, gtma-20). Immunoprecipitated proteins were spotted onto nitrocellulose membranes (Amersham, 10600007) and air dried for 1 hour. The membranes were blocked with 5% BSA in PBS for 1 hour and either incubated with 10 nM PAR chains overnight at 4˚C with or without the addition of 1% SDS or with 5 nM PAR chains for 2 hours at RT with or without the addition of 1% SDS. The membranes were washed extensively and incubated overnight with anti-PAR antibody (Trevigen, 4335-MC-100). Immunoblot analysis was performed to detect PAR and GFP expression.

**Co-immunoprecipitation**

GFP-tagged proteins (4 µg each) were electroporated as described above into approximately 7.5 million U2OS cells. Approximately 16 hours later, the cells were lysed in 250 µL lysis buffer (50 mM Tris pH 7.5, 1 mM EDTA, 150 mM NaCl, 1% Triton X-100, 1x Protease Inhibitor Cocktail Set III, 1x PMSF, 1x phosphatase inhibitor) and kept on ice for 5 minutes. Next, 12.5 µL of 5 M NaCl was added to the cell lysis buffer solution and kept on ice for 5 minutes. Cell lysis solution was centrifuge at max speed 4˚C for 12 minutes and the supernatant was moved to new pre-chilled tubes and kept on ice. Protein (500 µg) was diluted in 0.5X lysis buffer supplemented with 25 µL/mL of 5 M NaCl and 0.5x Protease Inhibitor Cocktail Set III, 1x PMSF, 1x phosphatase inhibitor). GFP antibody (1 µL/sample [Abcam, ab290]) was added and the sample were rotated end-over-end overnight at 4˚C. The following day, 50 µL of Protein G beads (Thermo, 88848) were pipetted in a tube (50 µL of beads/sample). The beads were pelleted with a magnet, the supernatant was removed, and the beads were equilibrated in 500 µL of lysis buffer. The beads were then resuspended with lysis buffer in the original slurry volume and 50 µL/sample was added. The samples we rotated end-over-end for 2 hours at 4˚C. Beads were washed 4x with 400 µL lysis buffer for 5 minutes at 4˚C (end-over-end rotation). Beads were resuspended in 25 µL lysis buffer and 1x protein dye was added. Samples were used in immunoblotting as described above.

**Statistics**

The statistical tests are indicated in the respective figure legends. Error bars indicate mean ± standard deviation (SD) except for the polysome profiling graphs and laser microirradiation graphs in which data represent mean ± standard error of the mean (SEM). *P* values of $\leq 0.05$ were considered significant. Analyses were performed using GraphPad Prism (v.8.0). For analysis of western blots, the Shapiro-Wilk test was used to test for normal distribution. If normally distributed, data were compared using a one-way analysis of variance (ANOVA) with *p* values adjusted for multiple comparisons. If the values did not pass the normality test, the values were log-transformed, and those values were analyzed using a one-way ANOVA with *p* values adjusted for multiple comparisons.

**Human subjects research**

This investigation was conducted according to the Declaration of Helsinki Principles. Written informed consent was received from participants before inclusion in the study according to protocol H-17698 Genetic and Biological Determinants of Bone Marrow Failure approved by the Institutional Review Board for Baylor College of Medicine.

#### SUPPLEMENTAL TABLES AND FIGURES

##### Supplemental Table 1. DBA LCLs used in this study.

| **BMF Number** | **RP Gene Mutated** | **DNA Change** | **Protein Change** |
|---|---|---|---|
| BMF16 | *RPL5* | c.535C>T | p.Arg179* |
| BMF51 | *RPS19* | c.1-1G>A | N/A |
| BMF54 | *RPS19* | c.280C>T | p.R94* |
| BMF89 | *RPL5* | c.70C>T | p.Arg24* |

**Supplemental Table 2. Sequences of siRNAs used in this study.**

| Name | Provider | siRNA ID | Sense Sequence | Antisense Sequence |
|---|---|---|---|---|
| RPS19-1 | Ambion | s535016 | GAUGAGAACUGGUUCUACAtt | UGUAGAACCAGUUCUCAUCgt |
| RPS19-2 | Ambion | s12323 | GCCGCAAACUGACACCUCAtt | UGAGGUGUCAGUUUGCGGCcg |
| RPL5-1 | Ambion | s12151 | GACGAGAGGGUAAAACUGAtt | UCAGUUUUACCCUCUCGUCgt |
| RPL5-2 | Ambion | s12152 | GGAGGAGAUGUAUAAGAAAttt | UUUCUUAUACAUCUCCUCCat |
| RAD51 | Ambion | s531930 | AGAAGGAGCUAAUAAAUAUtt | AUAUUUAUUAGCUCCUUCUtt |
| LIG4 | Ambion | s8181 | CAUUGAACCUUGUAAUUCUtt | AGAAUUACAAGGUUCAAUGta |
| TP53 | Horizon | J-003329-15 | Multiple targets. See providers website for accession numbers | |

**Supplemental Table 3. Antibodies used in this study.**

| Name | Provider | Catalog # | Species | Dilution |
|---|---|---|---|---|
| ATM | Abcam | Ab32420 | Rabbit | 1:5000 |
| ATR | Cell Sig | 13934 | Rabbit | 1:1000 |
| BRCA2 | Cell Sig | 10741 | Rabbit | 1:500 |
| DDX11 | Novus | NBP2-92510 | Rabbit | 1:1000 |
| DNA Ligase III (LIG3) | Thermo | PA5-21480 | Rabbit | 1:1000 |
| DNA Ligase IV (LIG4) | Abcam | Ab193353 | Rabbit | 1:1000 |
| DNA-PKcs | Abcam | Ab32566 | Rabbit | 1:1000 |
| ERCC1 | Novus | 34480 | Mouse | 1:665 |
| FLAG | Sigma | F7425 | Rabbit | 1:1000 |
| FLAG | Sigma | F3165 | Mouse | 1:5000 |
| GFP | Abcam | Ab290 | Rabbit | 1:1000 |
| Ku70 | Lab Vision | MS-329-P1 | Mouse | 1:500 |
| Ku86 | Sigma | Sc-5280 | Mouse | 1:200 |
| MRE11 | Cell Sig | 8344T | Rabbit | 1:1000 |
| MYC | Sigma | M4439 | Mouse | 1:5000 |
| NBS1 | Cell Sig | 8344T | Rabbit | 1:1000 |
| PAR | Trevigen | 4335-MC-100 | Mouse | 1:1000 |
| PARP1 | Bethyl | A301-376A | Rabbit | 1:500 |
| PARP2 | Active Motif | 39743 | Rabbit | 1:500 |
| PolQ | Sigma | SAB1402530 | Mouse | 1:1000 |
| p53 | Santa Cruz | SC-126 | Mouse | 1:200 |
| RAD51 | Abcam | Ab176458 | Rabbit | 1:1000 |
| RAD52 | Novus | 58116 | Rabbit | 1:500 |
| RPA2 | Invitrogen | PA5-22256 | Rabbit | 1:1000 |
| RPL5 | Abcam | Ab86863 | Rabbit | 1:500 |
| RPS19 | Bethyl | A304-002A | Rabbit | 1:1000 |
| XLF | Abcam | Ab33499 | Rabbit | 1:200 |
| XPF | Novus | 58407 | Rabbit | 1:1000 |
| XRCC1 | Abcam | Ab134056 | Rabbit | 1:1000 |
| XRCC4 | Thermo | PA5-82264 | Rabbit | 1:1000 |
| β-actin | Sigma | A5441 | Mouse | 1:5000 |
| β-tubulin | Abcam | ab6046 | Rabbit | 1:5000 |
| γ-H2AX | Sigma | 05-636 | Mouse | 1:1000 |
| IgG HRP-linked secondary antibody | Cell Sig | 7074 | Rabbit | 1:10000 |
| 680RD IgG secondary antibody | LI-COR | 926-68070 | Mouse | 1:5000 |
| 800CW IgG secondary antibody | LI-COR | 926-32211 | Rabbit | 1:5000 |

**Supplemental Table 4. RT-qPCR primers used in this study.**

| Target Gene | Forward Primer | Reverse Primer |
|---|---|---|
| *BRCA2* | ATCAGCTGGCTTCAACTCCA | TGGACAGGAAACATCATCTGC |
| *GAPDH* | CACATGGCCTCCAAGGAGTAAG | TACATGACAAGGTGCGGCTCCC |
| *LIG4* | CAGATATTGAGCACATTGAGAAG | GATCAGTGTAGTTATATCCATTTCG |
| *Luciferase* | ACACCCGAGGGGGATGATAA | CCAGATCCACAACCTTCGCT |
| *PARP1* | AAGAAATGCAGCGAGAGCAT | TCAGAGAACCCATCCACCTC |
| *RAD51* | GGCAATGCAGATGCAGCTTGAAGC | CCGTGAAATGGGTTGTGGGCCA |
| *XRCC4* | GGTTGGCTTCAGCTGCTGTAA | CTGATTCTCCTGAGGAGCCAT |

SUPP FIGURE 1
TO FIGURE 1

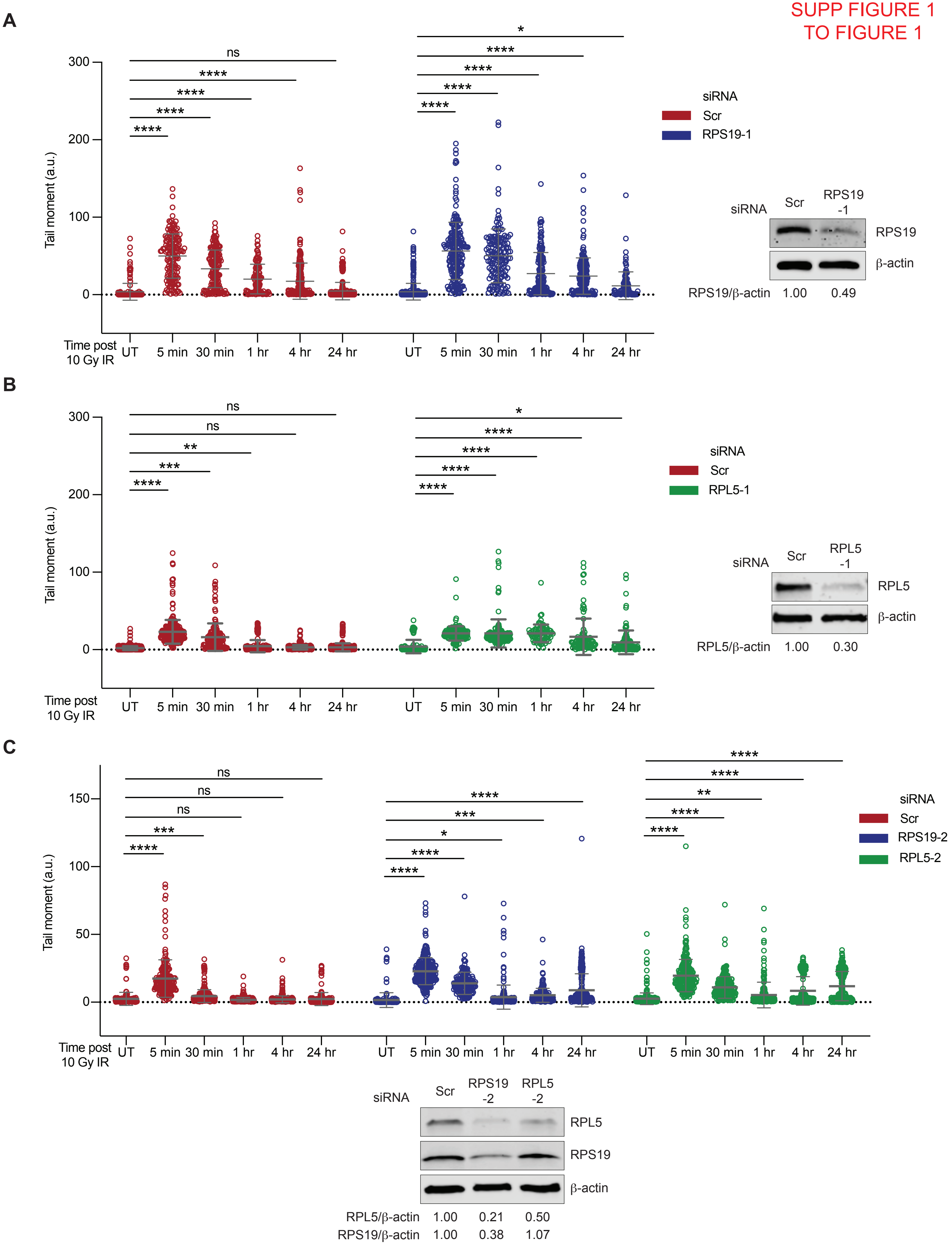

**Supplemental Figure 1.** (A) Neutral comet assay of siRNA KD RPS19-1 or scrambled (Scr) CD34+ cells UT or treated with 10 Gy IR and analyzed at the indicated time points post IR. Representative western blot showing KD of RPS19 using siRPS19-1 located to the right of graph. β-actin used as loading control. (B) Neutral comet assay of siRNA KD RPL5-1 or Scr CD34+ cells UT or treated with 10 Gy IR and analyzed at the indicated time points post IR. Representative western blot showing KD of RPL5 using siRPL5-1 located to the right of graph. β-actin used as loading control. (C) Neutral comet assay of siRNA KD RPS19-2, RPL5-2, or Scr CD34+ cells UT or treated with 10 Gy IR and analyzed at the indicated time points post IR. Representative western blot showing KD of RPS19 and RPL5 using siRPS19-2 and siRPL5-2, respectively, located below graph. β-actin used as loading control. Data represent mean ± SD compared using 1-way ANOVA with Dunnett's multiple comparisons test. ns not significant; *$P < 0.05$; **$P < 0.01$; ***$P < 0.001$; ****$P < 0.0001$.

SUPP FIGURE 2
TO FIGURE 2

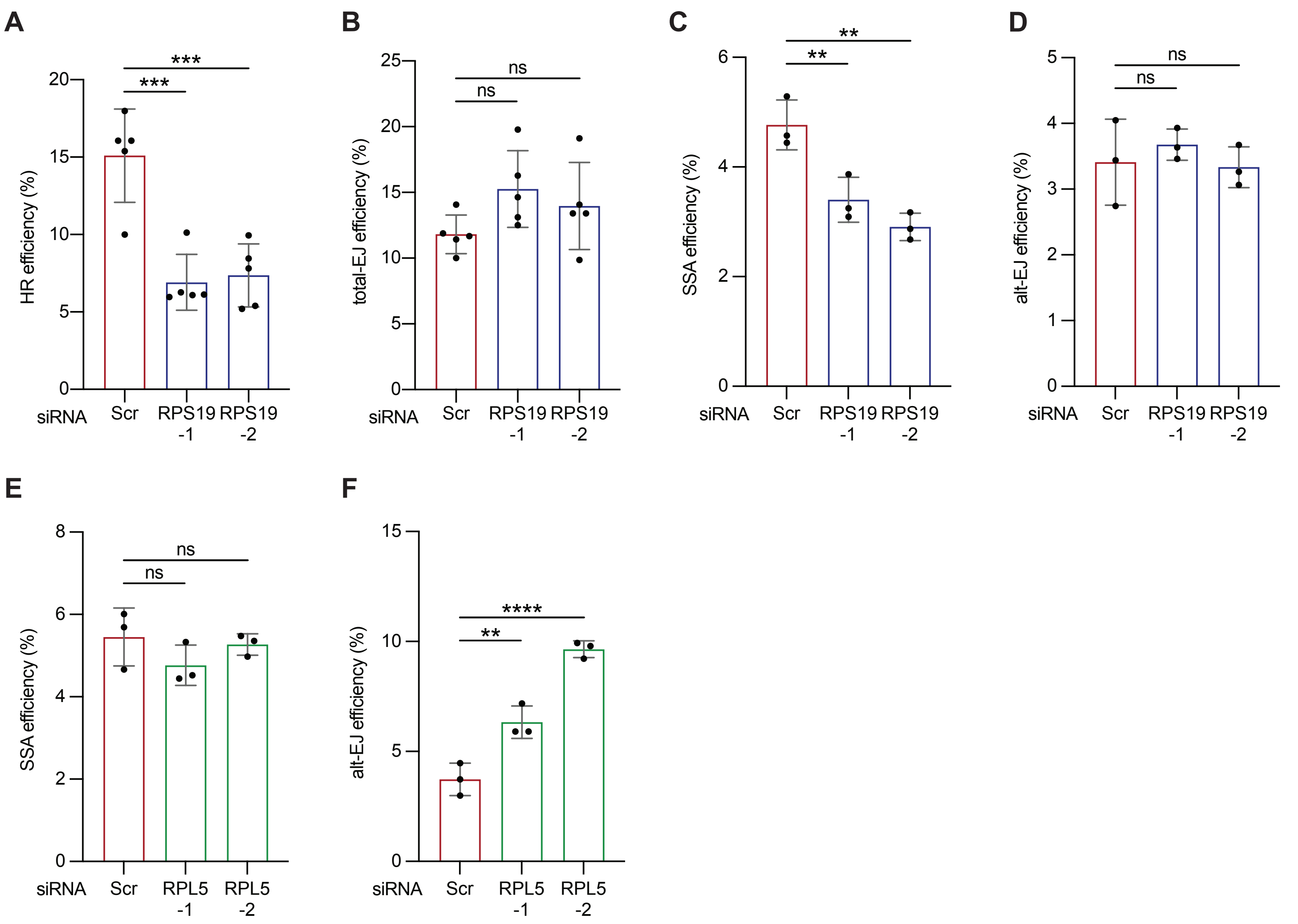


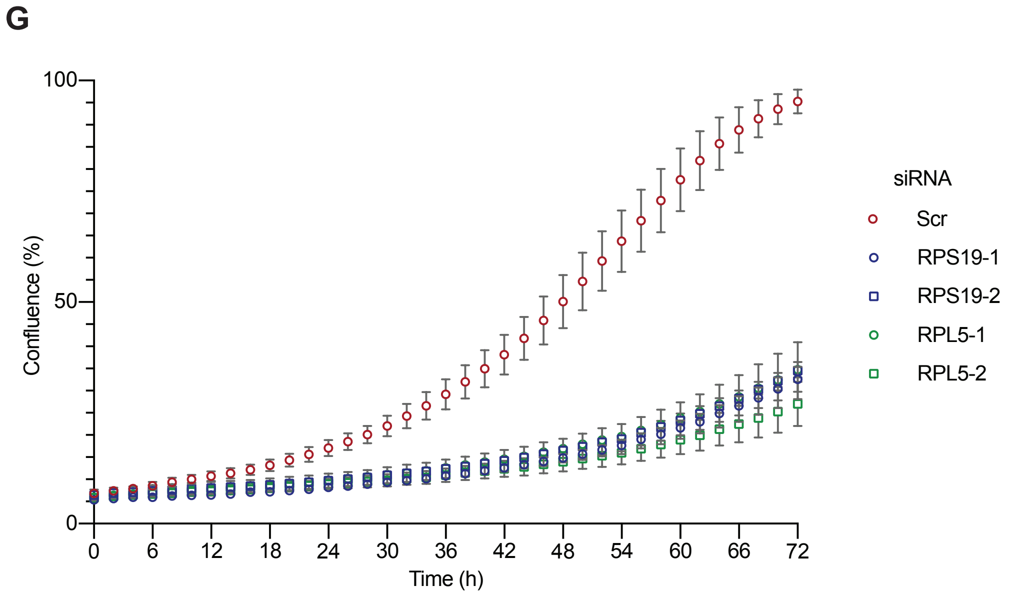


One-way ANOVA of proliferation experiment

| | |
|---|---|
| Scr vs. RPS19-1 | **** |
| Scr vs. RPS19-2 | **** |
| Scr vs. RPL5-1 | **** |
| Scr vs. RPL5-2 | **** |
| RPS19-1 vs. RPL5-1 | ns |
| RPS19-1 vs. RPL5-2 | ns |
| RPS19-2 vs. RPL5-1 | ns |
| RPS19-2 vs. RPL5-2 | ns |
| RPS19-1 vs. RPS19-2 | ns |
| RPL5-1 vs. RPL5-2 | ns |

**Supplemental Figure 2.** (A) HCT116 cells bearing an integrated DR-GFP HR reporter[1] transfected with Scr or one of two RPS19 siRNAs. After 48 hours, cells were co-transfected with an I-Sce1-expressing plasmid to induce a DSB in the DR-GFP reporter and a mCherry-expressing plasmid as a transfection control. Forty-eight hours later, cells were analyzed by flow cytometry. The repair efficiency was calculated by the proportion of GFP+ to mCherry+ cells (B) The same experiment in A except using HCT116 cells with an integrated EJ5-GFP total-EJ reporter.[2] (C) The same experiment in A except using U2OS cells with an integrated SA-GFP SSA reporter.[3] (D) The same experiment in A except using U2OS cells with an integrated EJ2-GFP alt-EJ reporter.[2] (E) The same experiment in C except using two different RPL5 siRNAs. (F) The same experiment in D except using two different RPL5 siRNAs. (G) Confluence of RPS19- and RPL5-KD cells assessed by live cell imaging over 96 hours. Each sample was performed in triplicate per experiment and the experiment was performed three times. Data in A–F represent three independent experiments. Data represent mean ± SD compared using 1-way ANOVA with Dunnett's multiple comparisons test. ns not significant; $^{**}P < 0.01$; $^{***}P < 0.001$; $^{****}P < 0.0001$.

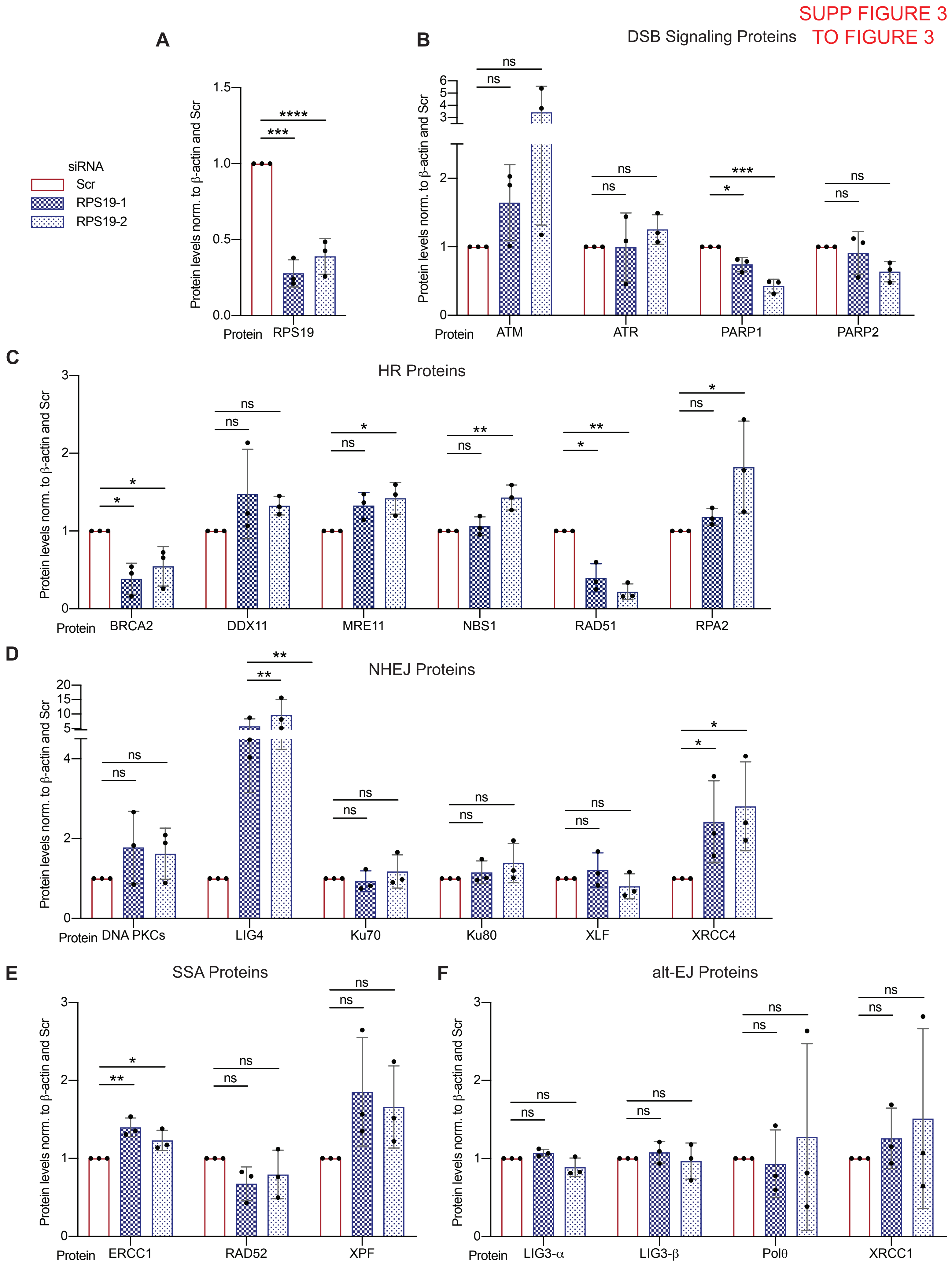
SUPP FIGURE 3
TO FIGURE 3
A
siRNA
Scr
RPS19-1
RPS19-2
Protein levels norm. to β-actin and Scr
Protein
RPS19
B
DSB Signaling Proteins
ATM
ATR
PARP1
PARP2
C
HR Proteins
BRCA2
DDX11
MRE11
NBS1
RAD51
RPA2
D
NHEJ Proteins
DNA PKCs
LIG4
Ku70
Ku80
XLF
XRCC4
E
SSA Proteins
ERCC1
RAD52
XPF
F
alt-EJ Proteins
LIG3-α
LIG3-β
Polθ
XRCC1

**Supplemental Figure 3.** All data represent protein quantification of RPS19-KD western blots as in Figure 3A. (A) Quantification of RPS19 protein levels in RPS19-KD cells using two different siRNAs normalized to β-actin and Scr. Data represent biological replicates. Representative images in Figure 4A. (B) Same in A except for the quantification of DSB signaling protein levels, (C) HR protein levels, (D) NHEJ protein levels, (E) SSA protein levels, and (F) alt-EJ protein levels. Data represent three independent experiments. Data represent mean ± SD. Shapiro-Wilk test was used to assess normality. The data were log-transformed and reanalyzed if samples did not pass the normality test. ns not significant; **P* < 0.05; ***P* < 0.01; ****P* < 0.001; *****P* < 0.0001.

SUPP FIGURE 4
TO FIGURE 3

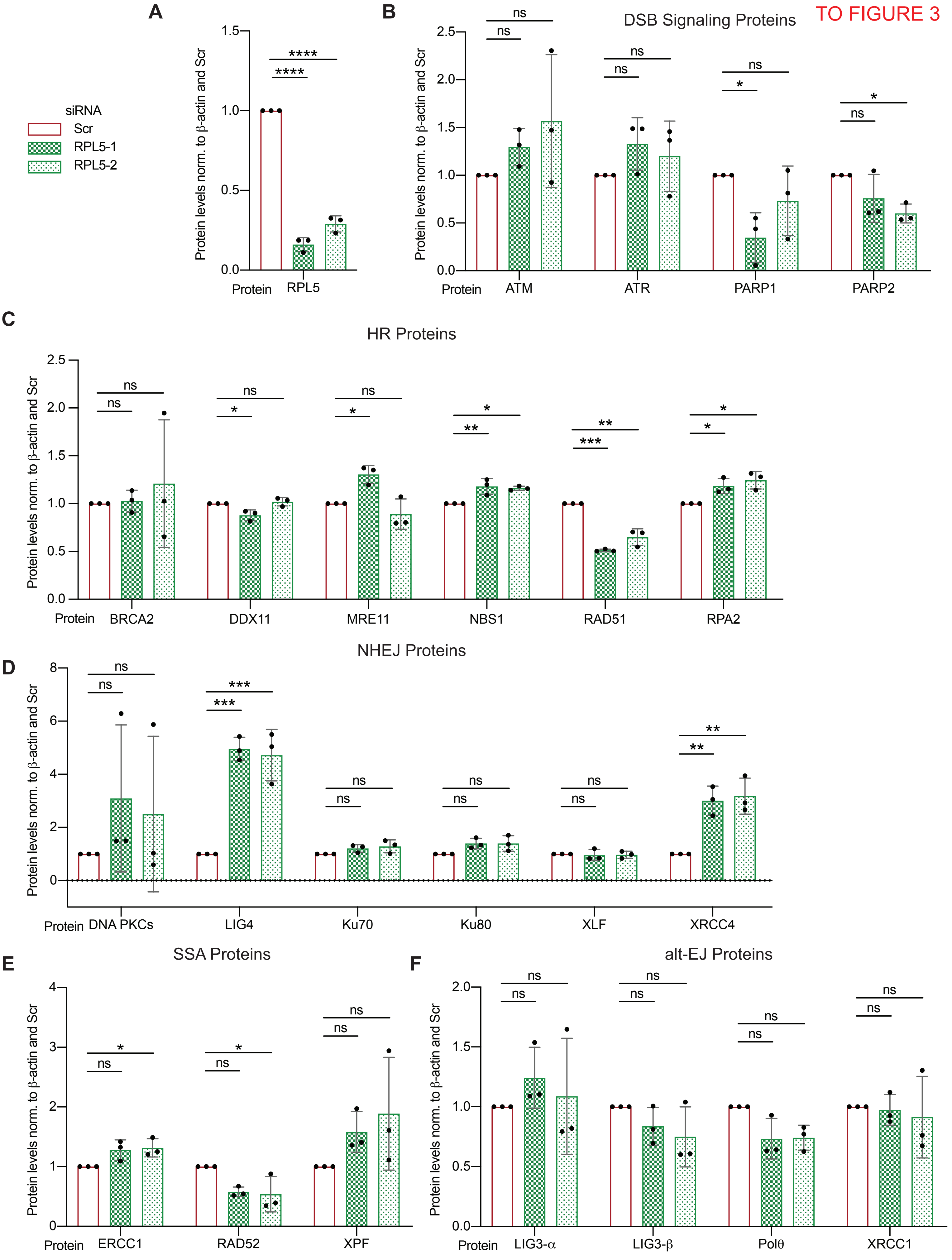

**Supplemental Figure 4.** All data represent protein quantification of RPL5-KD western blots as in Figure 3B. (A) Quantification of RPL5 protein levels in RPL5-KD cells using two different siRNAs normalized to β-actin and Scr. Data represent biological replicates. Representative images in Figure 4B. (B) Same in A except for the quantification of DSB signaling protein levels, (C) HR protein levels, (D) NHEJ protein levels, (E) SSA protein levels, and (F) alt-EJ protein levels. Data represent three independent experiments. Data represent mean ± SD. Shapiro-Wilk test was used to assess normality. The data were log-transformed and reanalyzed if samples did not pass the normality test. ns not significant; $*P < 0.05$; $**P < 0.01$; $***P < 0.001$; $****P < 0.0001$.

SUPP FIGURE 5
TO FIGURE 3

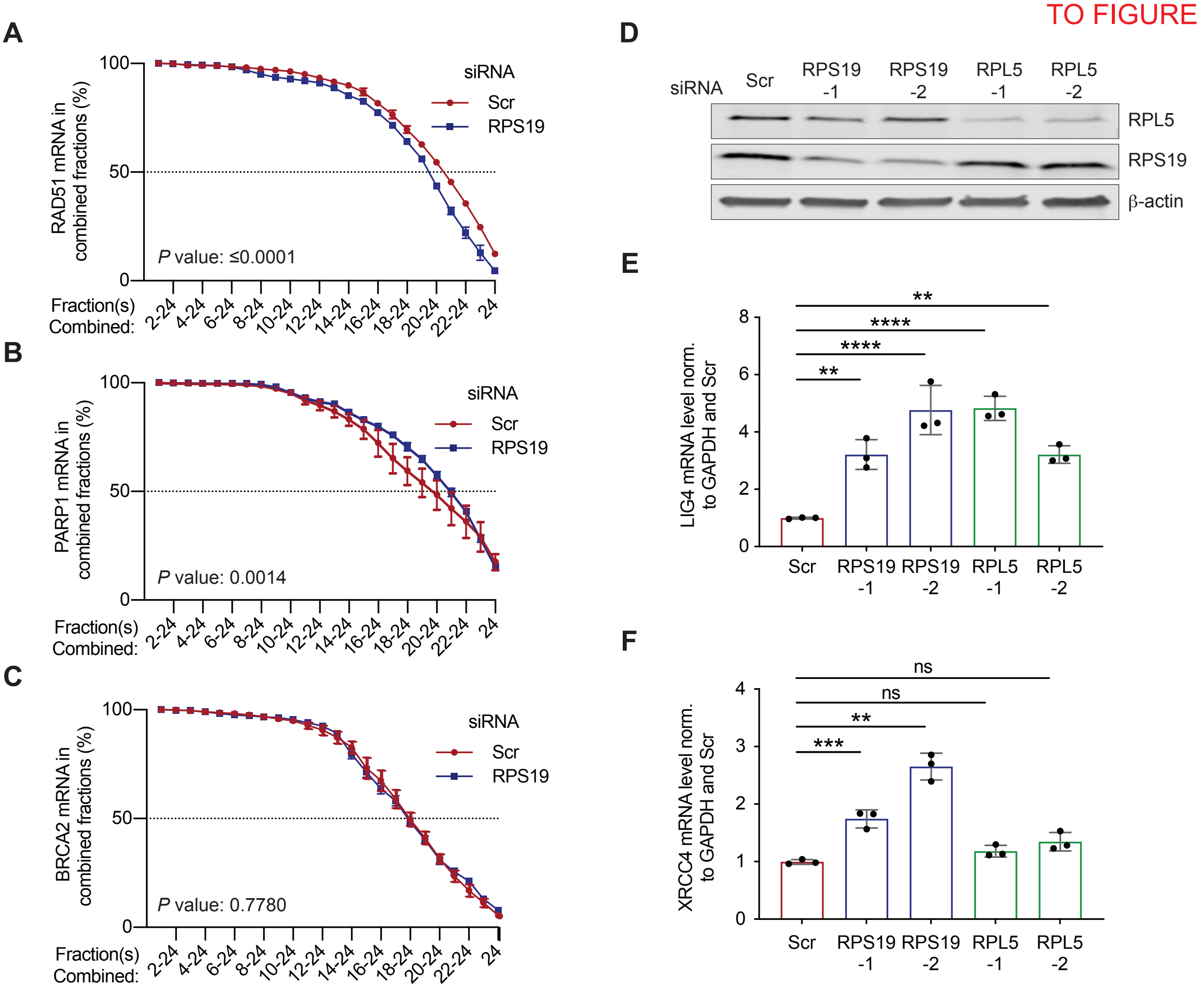


**Supplemental Figure 5.** (A) Cumulative RAD51 mRNA across combined fractions in Scr and RPS19-KD U2OS cells. (B) Same as A except assessing PARP1 mRNAs and (C) BRCA2 mRNAs. (D) Western blot of lysates prepared from RPS19- and RPL5-KD cells used for RT-qPCR experiments. β-actin used as a loading control. (E) Quantification of LIG4 mRNA levels in RPS19- and RPL5-KD U2OS cells normalized to GAPDH and Scr. (F) Same as E except quantification of XRCC4 mRNA levels. Data in A–C and E–F represent three biological replicates. Data in E–F represent mean ± SD compared using 1-way ANOVA with Dunnett's multiple comparisons test. ns not significant; ***P* < 0.01; ****P* < 0.001; *****P* < 0.0001.

SUPP FIGURE 6
TO FIGURE 3

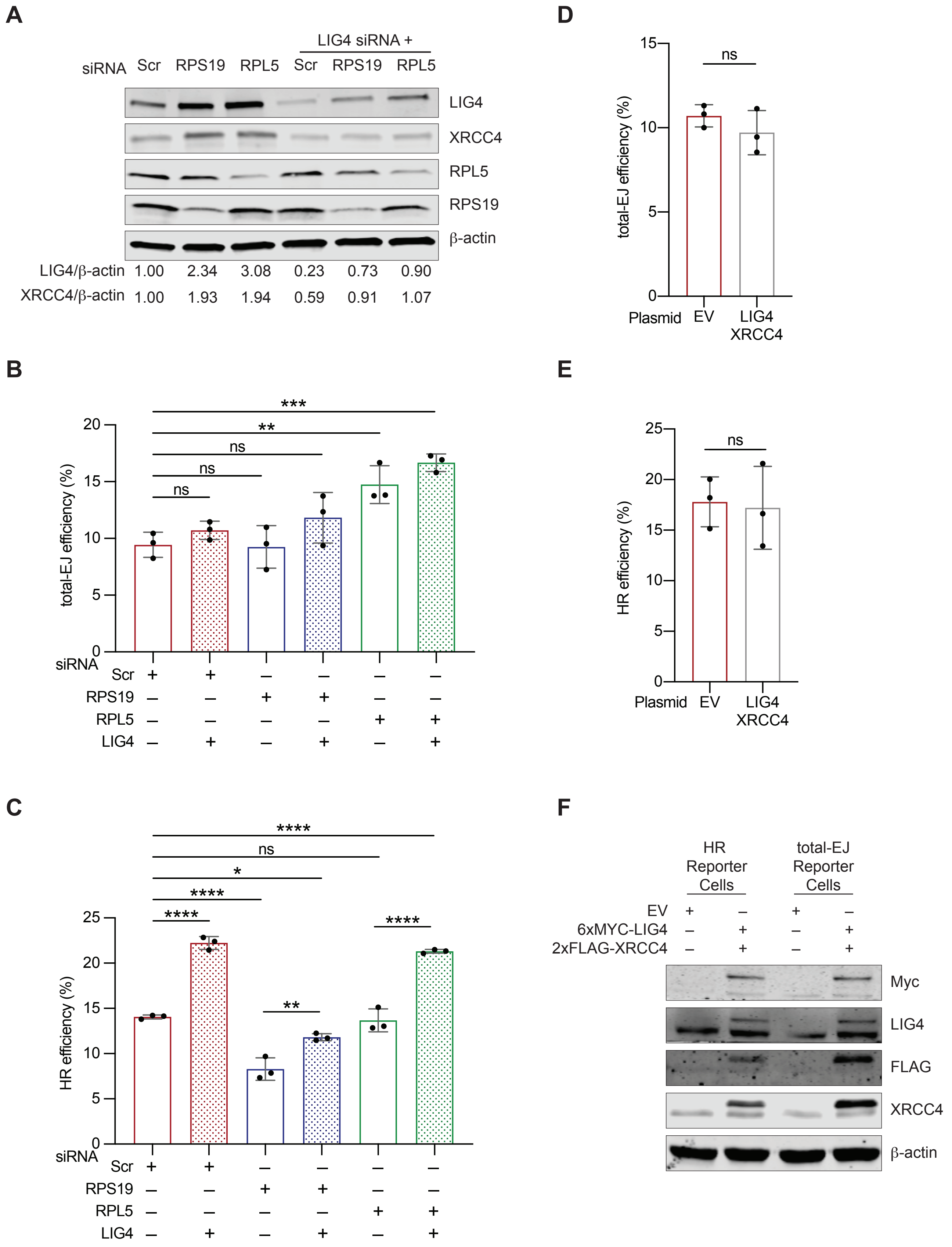

**Supplemental Figure 6.** (A) Western blot of RPS19- and RPL5-KD U2OS cells +/- LIG4-KD. β-actin used as a loading control. (B) U2OS cells bearing an integrated EJ5-GFP total-EJ reporter[2] transfected with Scr, RPS19, or RPL5 siRNAs. After 24 hours, cells were treated with LIG4-siRNA. Twenty-four hours later, cells were co-transfected with an I-Sce1-expressing plasmid to induce a DSB in the EJ5-GFP reporter and a mCherry-expressing plasmid as a transfection control. Forty-eight hours later, cells were analyzed by flow cytometry. The repair efficiency was calculated by the proportion of GFP+ to mCherry+ cells. (C) Same as B except using the DR-GFP HR reporter.[1] (D) Same as B except instead of KD, LIG4 and XRCC4 were co-transfected 48 hours prior to the co-transfection of the I-Sce1 and mCherry plasmids. (E) Same as C except using the DR-GFP HR reporter. (F) Western blot of lysates prepared from U2OS reporter cells overexpressing LIG4 and XRCC4. β-actin used as a loading control. Data in B–E represent three biological replicates. Data in B–C represent mean ± SD compared using 1-way ANOVA with Dunnett's multiple comparisons test. Data in D–E represent mean ± SD compared using unpaired t-tests. ns not significant; **P* < 0.05; ***P* < 0.01; ****P* < 0.001; *****P* < 0.0001.

SUPP FIGURE 7
TO FIGURE 5

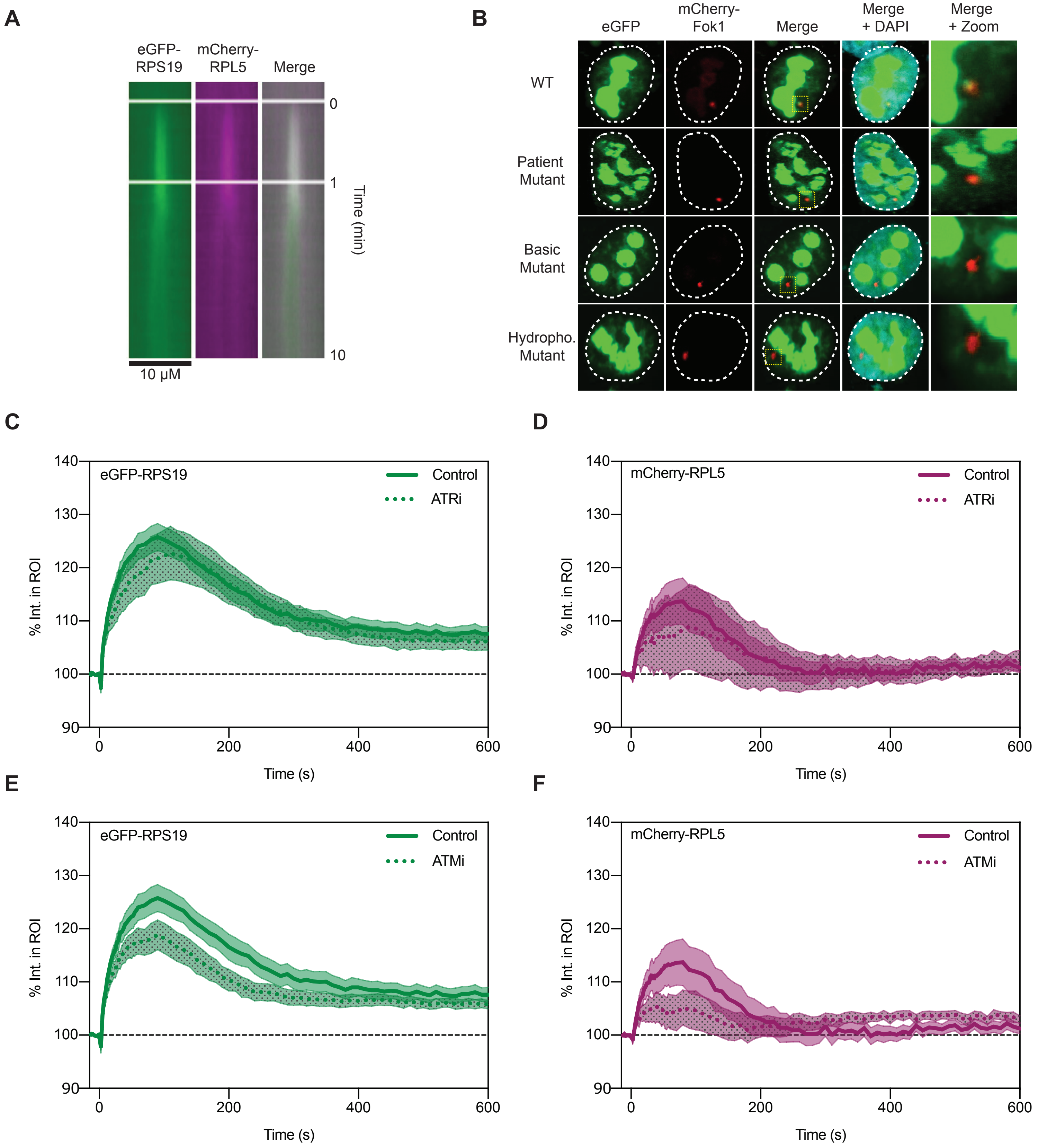


**Supplemental Figure 7.** (A) Kymograph of eGFP-RPS19 and mCherry-RPL5 spatial distribution in U2OS cells over 10 min. (B) Representative images of eGFP-RPS19 WT and mutants localization to mCherry-LacI-Fok1 DSB sites in U2OS cells. (C) Quantification of percent fluorescent intensity within the ROI of eGFP-RPS19 transfected U2OS cells treated with ATR (n = 8) inhibitor. (D) Same as C except cells were transfected with mCherry-RPL5 (n = 8). (E) Same as C except the cells treated with ATM (n = 8) inhibitor. (F) Same as D except the cells treated were with an ATM (n = 8) inhibitor. Controls in C–F are the same as in Figure 6A. Data in A–D represent mean ± SEM.

### SUPP FIGURE 8
### TO FIGURE 7

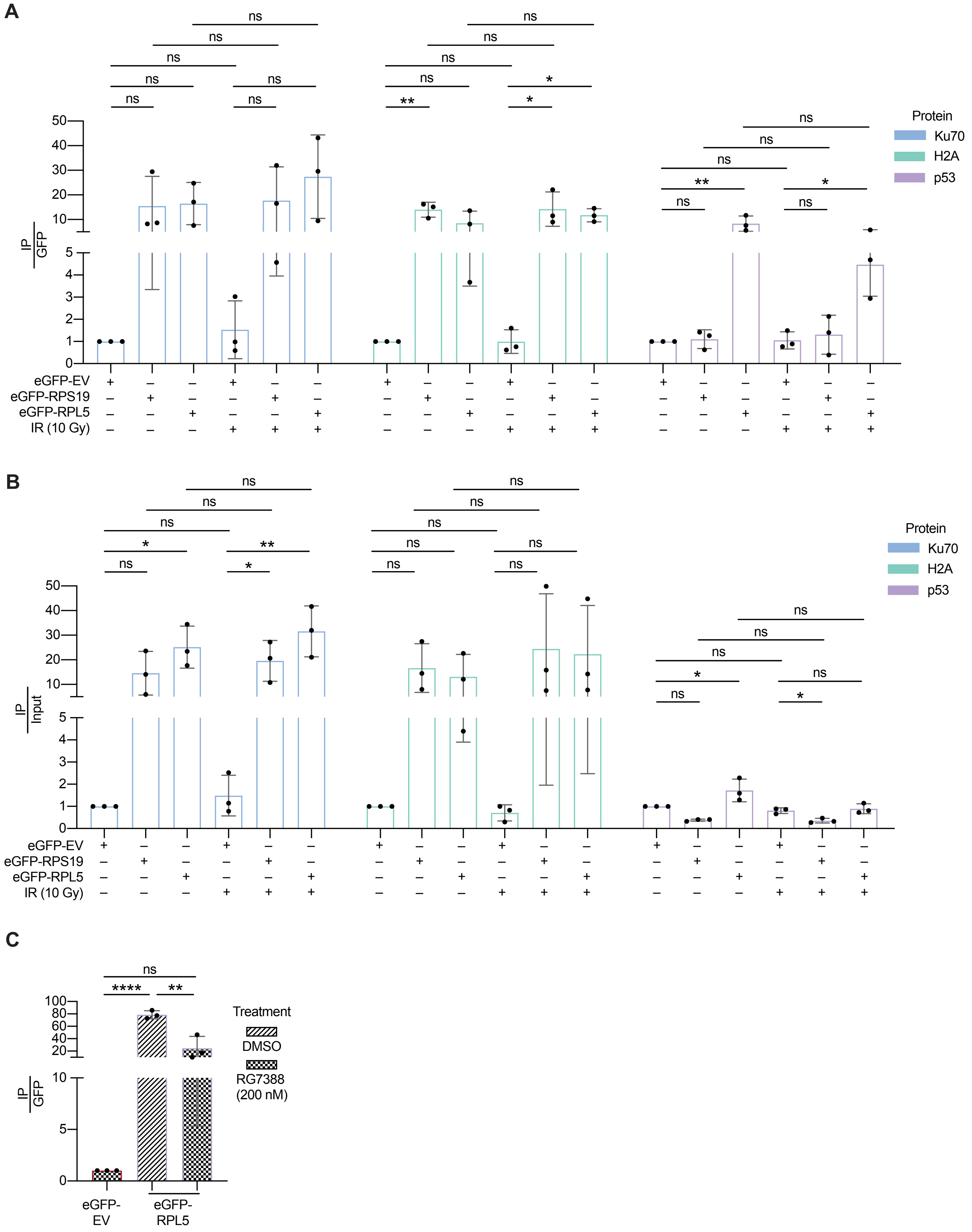

**Supplemental Figure 8.** (A) Quantification of proteins immunoprecipitated with eGFP-RPS19 and eGFP-RPL5 relative to GFP. (B) Same as A except relative to input. (C) Quantification of Figure 7B relative to GFP. Data represent mean ± SD compared using unpaired t-tests of three biological replicates. ns not significant; *$P < 0.05$; **$P < 0.01$; ****$P < 0.0001$.

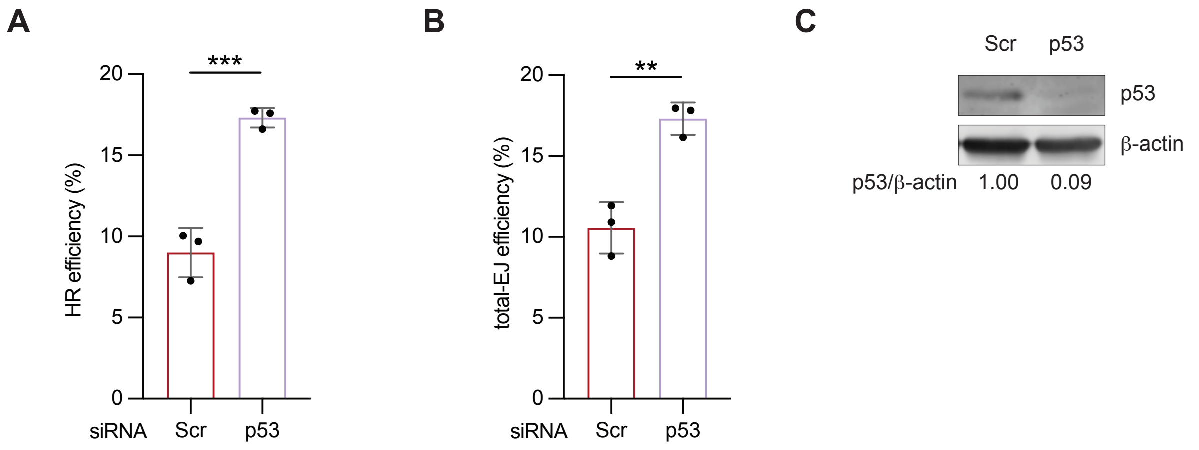


**Supplemental Figure 9.** (A) U2OS cells bearing an integrated DR-GFP HR reporter[1] transfected with Scr or p53 siRNA. After 48 hours, cells were co-transfected with an I-SceI-expressing plasmid to induce a DSB in the DR-GFP reporter and a mCherry-expressing plasmid as a transfection control. Forty-eight hours later, cells were analyzed by flow cytometry. The repair efficiency was calculated by the proportion of GFP+ to mCherry+ cells. (B) The same experiment in A except using U2OS cells with an integrated EJ5-GFP total-EJ reporter. (C) Representative western blot of p53 protein levels in p53-KD cells. β-actin used as a loading control. Data in A–B represent mean ± SD compared using unpaired t-tests of three biological replicates. $^{**}P < 0.01$; $^{***}P < 0.001$.
